# A mouse model of ZTTK syndrome reveals indispensable SON functions in organ development and hematopoiesis

**DOI:** 10.1101/2023.11.19.567732

**Authors:** Lana Vukadin, Bohye Park, Mostafa Mohamed, Huashi Li, Amr Elkholy, Alex Torrelli-Diljohn, Jung-Hyun Kim, Kyuho Jeong, James M Murphy, Caitlin A. Harvey, Sophia Dunlap, Leah Gehrs, Hanna Lee, Hyung-Gyoon Kim, Seth N. Lee, Denise Stanford, Robert A. Barrington, Jeremy B. Foote, Anna G. Sorace, Robert S. Welner, Blake E. Hildreth, Ssang-Taek Steve Lim, Eun-Young Erin Ahn

**Affiliations:** Department of Pathology, Division of Molecular and Cellular Pathology, University of Alabama at Birmingham, Birmingham, AL, USA; Department of Neurosurgery, University of Alabama at Birmingham, Birmingham, AL, USA; Metastasis Branch, Division of Cancer Biology, National Cancer Center, Goyang, Gyeonggi-do, Korea; Department of Medicine, College of Medicine, Dongguk University, Gyeongju, Korea; Department of Radiology, University of Alabama at Birmingham, Birmingham, AL, USA; Department of Medicine, Cystic Fibrosis Research Center, University of Alabama at Birmingham, Birmingham, AL, USA; Department of Microbiology and Immunology, College of Medicine, University of South Alabama, Mobile, AL, USA; Department of Microbiology, University of Alabama at Birmingham, Birmingham, AL, USA; O’Neal Comprehensive Cancer Center, University of Alabama at Birmingham, Birmingham, AL, USA; Department of Medicine, Division of Hematology and Oncology, University of Alabama at Birmingham, Birmingham, AL, USA

## Abstract

Rare diseases are underrepresented in biomedical research, leading to insufficient awareness. Zhu-Tokita-Takenouchi-Kim (ZTTK) syndrome is a rare disease caused by genetic alterations that result in heterozygous loss-of-function of SON. While ZTTK syndrome patients suffer from numerous symptoms, the lack of model organisms hamper our understanding of both SON and this complex syndrome. Here, we developed *Son* haploinsufficiency (*Son^+/−^*) mice as a model of ZTTK syndrome and identified the indispensable roles of *Son* in organ development and hematopoiesis. *Son^+/−^* mice recapitulated clinical symptoms of ZTTK syndrome, including growth retardation, cognitive impairment, skeletal abnormalities, and kidney agenesis. Furthermore, we identified hematopoietic abnormalities in *Son^+/−^* mice, similar to those observed in human patients. Surface marker analyses and single-cell transcriptome profiling of hematopoietic stem and progenitor cells revealed that *Son* haploinsufficiency inclines cell fate toward the myeloid lineage but compromises lymphoid lineage development by reducing key genes required for lymphoid and B cell lineage specification. Additionally, *Son* haploinsufficiency causes inappropriate activation of erythroid genes and impaired erythroid maturation. These findings highlight the importance of the full gene dosage of *Son* in organ development and hematopoiesis. Our model serves as an invaluable research tool for this rare disease and related disorders associated with SON dysfunction.

## Introduction

Rare diseases are clinical conditions with a low prevalence, affecting fewer than 200,000 people in the United States. However, there are more than 7,000 different types of rare diseases, and over 350 million individuals suffer from rare diseases globally, with more than half of them being children (1–3). Nevertheless, most rare diseases are set aside in a “dark zone” where attention from the medical and research community is lacking. While advancements in whole exome and genome sequencing are rapidly unveiling previously undiagnosed rare genetic diseases, further research following the initial identification of these diseases is not sufficient to move forward (1, 4). This is partially due to a lack of proper model organisms that faithfully recapitulate the clinical features of human patients (5). Establishing proper animal models for newly identified human genetic diseases is critical to provide patients, families, and clinicians with further clinical characterization and to develop potential therapies. This effort is particularly critical to define clinical features of rare diseases, which are often ambiguous due to the small patient number.

Zhu-Tokita-Takenouchi-Kim (ZTTK) syndrome (also known as SON-related disorder) is a recently identified rare genetic disease characterized by developmental delay and multiple congenital anomalies (6–9). This syndrome is caused by mutations or entire/partial deletion of the *SON* gene from one allele, resulting in heterozygous loss-of-function (LoF) of *SON* (6, 7). After initial reports of 28 individuals with *SON* LoF variants in 2016, the number of cases continues to increase, revealing various clinical features that were not noticed previously. Although the initial effort focused on patients’ symptoms associated with developmental delay (DD) and intellectual disability (ID), further examination of the patients and subsequent research identified that *SON* LoF causes a wide spectrum of clinical features besides DD and ID (10–16). Thus, ZTTK syndrome is a complex, multi-system developmental disorder (OMIM #617140 and MedGen UID 934663).

Our initial discovery of this syndrome confirming the pathogenicity of SON LoF led to family connections and our continuous support for the families greatly contributed to the launch of an official non-profit organization, the ZTTK SON-Shine Foundation (https://zttksonshinefoundation.org/).

The *SON* gene is located on human chromosome 21 and encodes the SON protein, which possesses both DNA- and RNA-binding abilities and mainly localizes in nuclear speckles. Compromised SON function leads to aberrant and alternative RNA splicing, particularly for transcripts bearing weak splice sites (17–20), and SON-mediated RNA splicing is critical for maintaining pluripotency in embryonic stem cells (21). We have also shown that SON binds to DNA and suppresses H3K4me3 modifications at transcription start sites by interacting with Menin and sequestering it from the MLL1/2 methyltransferase complex (22). Recent attention to the role of nuclear speckles in gene expression has revealed that SON serves as the core of nuclear speckles (23) and enhances p53-mediated transcription (24). Therefore, SON governs the expression of a myriad of target genes by regulating transcription, RNA splicing, and nuclear speckle assembly, while exerting its effects on specific sets of genes.

To extend our understanding of the clinical features associated with ZTTK syndrome and to study how *Son* loss affects developmental processes and growth, we created mouse models with the floxed *Son* gene, as well as mice with a germline *Son* deletion. Here, we report on the indispensable roles of *Son* in embryo development and demonstrate that mice with heterozygous *Son* loss recapitulate multiple clinical features of human ZTTK syndrome. Importantly, our mouse model identified hematopoietic abnormalities, which were found in human ZTTK syndrome patients. Moreover, we conducted surface marker-based phenotypic analysis combined with single-cell transcriptome analysis to determine how *Son* haploinsufficiency affects hematopoietic lineage determination and differentiation by perturbing critical gene expression.

## Results

### Generation of *Son*-floxed mice (*Son^flox/flox^*) and constitutive *Son* knockout mice to establish a mouse model of human ZTTK syndrome

Given the absence of any reports on human cases with homozygous *Son* LoF variants, along with potential viability and fertility issues caused by heterozygous LoF of *Son*, we adopted a *Son*-floxed mouse model approach rather than a conventional knockout strategy. We inserted two loxP sites flanking the mouse *Son* exon 2 (Figure 1A), and the founder mice (F0) with one allele floxed by the loxP sites (*Son^flox/wt^*) were obtained. Then, the F0 mice were backcrossed to C57BL/6, and F1 heterozygous mice were confirmed by Southern blot and PCR (Supplemental Figure 1, A-C). Subsequently, we generated homozygous mice (*Son^flox/flox^*). The resulting *Son^flox/flox^* line will prove useful in studying tissue-specific and developmental stage/age-specific Son function.

**Figure 1.**
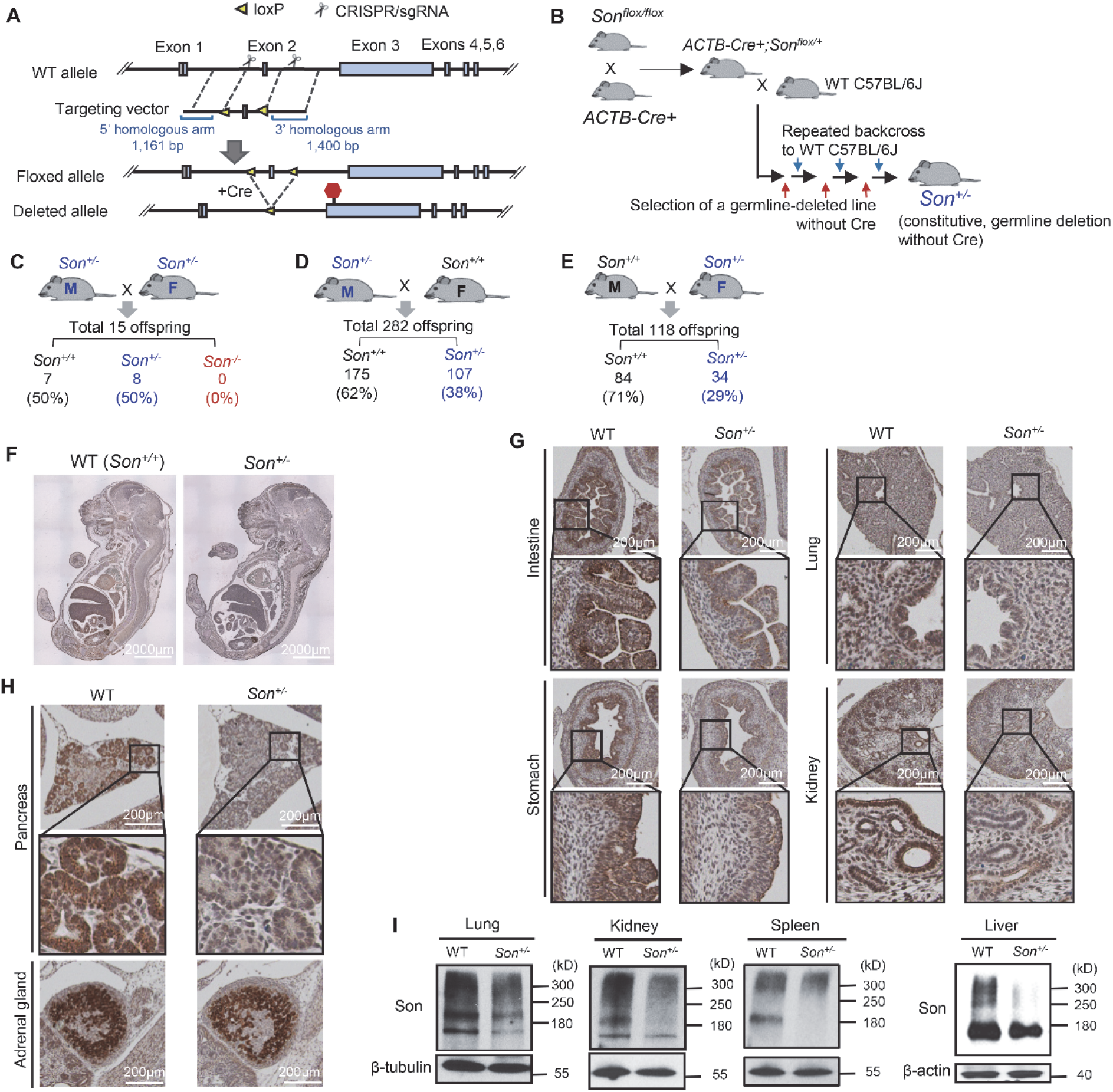
Generation of the *Son^+/−^*mouse models and characterization of Son expression patterns in embryos and organs. (**A**) Schematic diagram of the targeted locus and the strategy to insert loxP sites flanking *Son* exon 2. (**B**) A schematic illustrating the strategy to generate *Son^+/−^* mouse with permanent, germline deletion of the floxed sequence. **(C)** *Son^+/−^* intercrossing revealed extremely low fertility of this breeding scheme and embryonic lethality of *Son^−/−^* . (**D**) Breeding strategy crossing male *Son^+/−^* with female WT mice, showing the less *Son^+/−^* offspring than expected by the Mendelian ratio (**E**) Breeding strategy crossing male WT with female *Son^+/−^* mice, showing further decreased *Son^+/−^* offspring. (**F**) IHC staining of Son in sagittal sections of WT and *Son^+/−^* embryos (E16). (**G** and **H**) IHC staining of Son in E16 embryonic organs with enlarged images of the boxed region. (**I**) Expression of the Son protein in WT and *Son^+/−^* mice was analyzed by Western blot. β-tubulin or β-actin was used as a loading control. WB data are representative of n=3 independent experiments.

To create a mouse model that mimics human ZTTK syndrome, we proceeded to generate mice with heterozygous *Son* deletion in the entire body by crossing *Son^flox/flox^* mice with *β-actin-cre* (*ACTB-cre*) mice to ubiquitously delete the floxed sequence. After confirming that the *ACTB-Cre^+^*; *Son ^flox/wt^* mice were viable, we backcrossed them with wild-type C57BL/6J mice multiple times (>6 generations). Through this process, we selected mice with germline deletion of the floxed sequence in the *Son* gene, without Cre recombinase expression, to establish the line with constitutive, heterozygous deletion of *Son* (*Son^+/−^*) from germline (Figure 1B and Supplemental Figure 1, D and E). Deletion of the floxed region resulted in a frameshift due to the loss of exon 2 and a subsequent premature termination codon in exon 3.

### Homozygous deletion of *Son* (*Son^−/−^*) results in embryonic lethality occurring at an embryonic stage earlier than E6.0

To investigate the effects of heterozygous (*Son^+/−^*) and homozygous knockout (*Son^−/−^*) of *Son* on embryo viability and development, we performed a cross between a *Son^+/−^* male and a *Son^+/−^* female. However, this *Son^+/−^* intercrossing did not yield any *Son^−/−^* offspring (Figure 1C), indicating that constitutive, homozygous *Son* deletion (*Son* knockout; KO) causes embryonic lethality during development. To identify the embryonic stage at which lethality occurs, we collected numerous embryos at different stages of development and genotyped them by PCR. Despite our attempts to genotype as early as E6.5-7.5 (gastrulation), we did not detect any embryos showing the *Son^−/−^* genotype. These results indicate that *Son* is indispensable for mouse embryo development, and *Son*-null embryos with constitutive homozygous *Son* deletion lead to lethality earlier than the gastrulation stage. This finding is consistent with the fact that no human ZTTK syndrome patients bearing homozygous *Son* loss-of-function mutations have been identified. Additionally, we observed that *Son^+/−^* intercrossing breeding cages exhibited extremely low levels of fertility, generating only 15 pups in total (7 *Son^+/+^* and 8 *Son^+/−^*) from more than 10 breeding pairs attempted over 2 years (Figure 1C).

### Mice with heterozygous deletion of *Son* (*Son^+/−^*) as a model for ZTTK syndrome were born and viable, but were not produced at a normal Mendelian ratio

From the *Son^+/−^* intercrossing described above, we found that the *Son^+/−^* mice were born and viable. However, the ratio of *Son^+/+^* and *Son^+/−^* offspring was around 1:1 (7/15 and 8/15), which is less than the expected Mendelian ratio (*Son^+/+^* : *Son^+/−^* ratio to be 1:2). We further crossed *Son^+/−^* mice with *Son^+/+^* wild-type (WT) mice and found that the frequency of *Son^+/−^* offspring was significantly lower than expected (Figure 1, D and E; the expected frequency of *Son^+/−^* is 50%). Interestingly, the frequency of *Son^+/−^* offspring was affected by whether the breeder with *Son^+/−^* genotype was male or female in the breeding pair. When a *Son^+/−^* male was crossed with a WT female, about 38% of the offspring were *Son^+/−^* (Figure 1D). However, when *Son^+/−^* females were crossed with a WT male, only 29% of the offspring were *Son^+/−^* (Figure 1E), and the breeding scheme using female *Son^+/−^* (Figure 1E) generated significantly fewer offspring than breeding using male *Son^+/−^*. These findings suggest that constitutive, heterozygous deletion of *Son* during embryonic development partially causes prenatal lethality, which may be further increased when the *Son*-deleted allele is passed from a female (from an oocyte) to the offspring. It is also possible that the compromised production and/or viability of oocytes contribute to the difficulty of getting offspring from *Son^+/−^* female mice. Further studies are needed to determine the exact cause and molecular basis of these findings.

### Son is ubiquitously expressed, with the highest levels in the epithelial lining of gastrointestinal tracts and the gland structures in developing embryos, and a general reduction in Son expression was confirmed in *Son^+/−^*embryos

Next, we performed H&E staining of sagittal sections of whole embryos (E16-17) to investigate the gross morphology of *Son^+/−^* embryos. We found no noticeable defects in the morphology of the *Son^+/−^* embryos compared to *Son^+/+^* (Supplemental Figure 2A). Immunohistochemistry (IHC) staining for Son confirmed that *Son^+/−^* embryos had reduced Son protein levels throughout the body compared to *Son^+/+^* WT embryos (Figure 1F). We found that while Son protein was ubiquitously expressed in the embryo, certain tissues/organs showed particularly high levels of expression. These included gastrointestinal epithelial cells such as intestinal epithelium crypts and villi (with higher levels in the crypts) and inner layers of the stomach, the lung (particularly in airway epithelial cells), and the kidney (tubular epithelial cells) (Figure 1G and Supplemental Figure 2B). We also found that organs with gland structures, such as pancreatic islets, thyroid gland, and adrenal glands, exhibited high levels of Son protein expression (Figure 1H and Supplemental Figure 2B). Relatively moderate to low levels of Son were detected in cartilage, circular muscle layers of gastrointestinal tracts, and the ventricular zone of the brain (Figure 1F).

Interestingly, the majority of tissues/organs with high Son expression levels were endoderm-derived epithelial linings and glands, while mesoderm- and ectoderm-derived tissues/organs showed lower levels of Son expression. The negative control without primary antibodies was completely devoid of brown staining, demonstrating the specificity of our staining (Supplemental Figure 2C). Our extensive IHC analyses confirmed the reduction of Son in *Son^+/−^* embryos and provided valuable information on the endogenous Son protein expression pattern in developing mouse embryos.

### Examining *Son^+/−^* mice revealed various patterns and sizes of the Son protein expressed among different tissues/organs

Previous studies, both from our group and others, have shown that multiple bands are detected in SON Western blot experiments, ranging from over 300 kDa to approximately 120 kDa, in several human cell lines as well as human peripheral blood mononuclear cells (PBMCs) (6, 17, 25, 26). It has been speculated that SON is present in multiple forms besides the one with a calculated molecular weight, as all of these bands are eliminated by SON siRNA (26). However, previous Western blots detected SON in mainly in cancer cell lines, and have not examined the protein expression pattern of SON in normal tissues/organs besides PBMCs. To take advantage of whole-body, constitutive *Son^+/−^* mice, we performed Western blotting on different tissues/organs isolated from these mice and compared the patterns with those from *Son^+/+^* WT mice (Figure 1I and Supplemental Figure 2D). While we detected bands around and above the 300 kDa molecular weight marker, which should represent the full-length Son protein, we also observed a band migrating to a position slightly below the 180 kDa marker. Interestingly, we detected different band patterns in different tissues/organs, and all bands showed decreased intensity in tissues/organs from *Son^+/−^* mice (Figure 1I). These suggest that all bands are *Son* gene products, not non-specific bands, representing potential protein products from different transcript variants or post-translationally modified forms.

### *Son^+/−^* mice show significant growth retardation and fail to gain weight properly, which resembles the symptom found in human ZTTK syndrome patients

Mild or severe growth retardation is a common phenotype observed in human ZTTK syndrome patients, with more than 50% of examined patients showing short stature and low weight (11). None of the infant-adolescent patients showed a growth parameter above the 97th percentile (11). We therefore monitored the growth of *Son^+/−^* mice and their WT (*Son^+/+^*) littermates, and found the significant growth retardation of *Son^+/−^* mice which persisted and worsened throughout their lifetime. Both male and female *Son^+/−^* mice gradually gained weight, but not as much as their WT. Notably, after 21 weeks of age, *Son^+/−^* mice barely gained body weight, with the average weights gained during the age of 21 to 60 weeks being only 3.3g for male and 2.5g for female. Meanwhile, male and female WT littermates gained 13.2g and 7.3g, respectively, during the same period. At 60 weeks of age, the body weights of male and female *Son^+/−^* mice were approximately only 58% of those of male and female WT littermates ((Figure 2, A and B, and Supplemental Figure 3 for measurement data).

**Figure 2.**
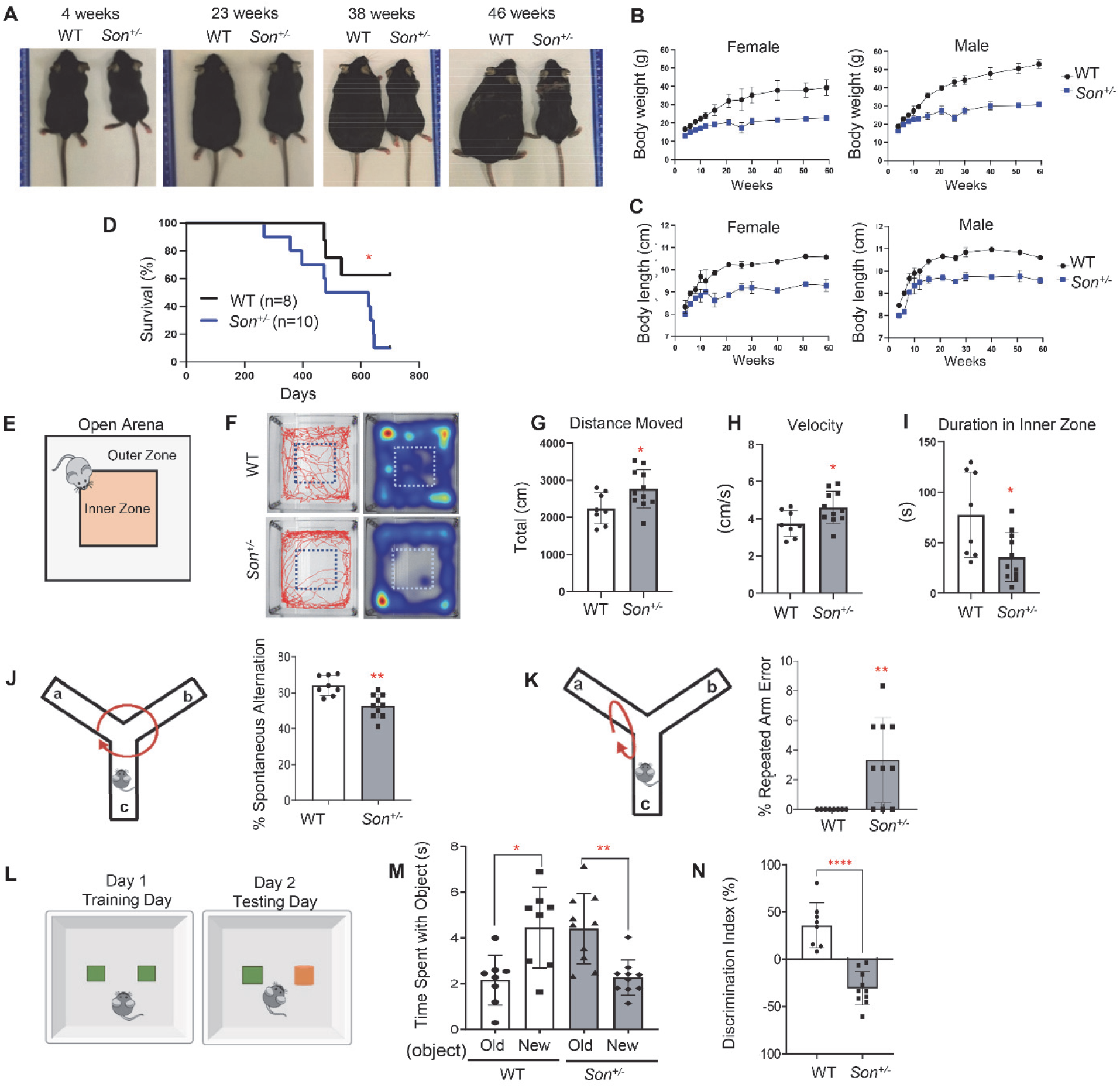
*Son^+/−^* mice show growth retardation, weight gain failure, and behavioral phenotypes, resembling clinical features of human ZTTK syndrome. (**A**) Representative images indicating body size differences between WT and *Son^+/−^* mice at various ages. (**B** and **C**) The body weight (**B**) and the body length (**C**) of female and male mice were measured at different ages up to 60 weeks. Data are presented as mean ±SD, n=5-12. (**D**) Kaplan-Meier curves of survival data in *Son^+/−^* mice compared to WT with a follow-up duration of 700 days of age. *p < 0.05. (**E**) Schematic depicting the open field test. **(F)** Representative tracks of WT and *Son^+/−^* showing the total distance traveled by the indicated mouse, and the representative heatmaps of time mouse spent in the open arena. (**G**) Total distance moved was calculated by measuring total centimeters (cm) traveled. (**H**) Velocity was calculated by measuring centimeters traveled per second (cm/s). (**I**) Time (seconds) spent in the Inner Zone of the open area. (**J** and **K**) Schematics and the graphs of percent level showing correct spontaneous alternation (**J**) and incorrect spontaneous alternation, also called repeated arm error (**K**) of WT and *Son^+/−^* mice in the Y-maze test. (**L**) Schematic depicting the experimental design of the Novel Object Recognition test. Old object, depicted in green; new object, depicted in orange. (**M**) Time spent with the old versus new object was calculated in seconds (s). (**N**) A discrimination index was calculated using the percentage of the ratio between the time spent (*T*) per exploration number (*N*) of the novel object by the time spent per exploration number of both objects (*T*new/*N*new – *T*old/*N*old / *T*new/*N*new + *T*old/*N*old). For graphs in (**G**), (**H**), (**I**), (**J**), (**K**), (**M**), and (**N**), data are presented as mean ±SD, n=8-10 per group. * p< 0.05 **p<0.01 ****p<0.0001

We also measured the body length of *Son^+/−^* mice and their WT littermates between the ages of 4 and 60 weeks. During this time, the body length of *Son^+/−^* mice and WT littermates was about 90-94% of WT body length from ages 4-12 weeks and remained 86-90% of WT body length between ages 12-60 weeks (Figure 2C and Supplemental Figure 3 for measurement data). These analyses demonstrated that *Son^+/−^* mice showed growth retardation, particularly a failure to gain weight, recapitulating the phenotype of human ZTTK syndrome patients.

Despite significant growth retardation and failure to gain weight, *Son^+/−^* mice did not show a noticeable mortality until 300-400 days. However, after this time point, they showed an increased death compared to their WT littermates when monitored up to 700 days, under standard care in the pathogen-free animal facility (Figure 2D).

### *Son^+/−^* mice show signs of hyperactivity, anxiety-like behavior, and cognitive impairment

Nearly 100% of individuals with ZTTK syndrome exhibit intellectual disability (6, 7, 11), and approximately 57% (26 out of 46) of those examined show behavior problems (11). The ZTTK SON-Shine foundation has reported deficits in spatial memory and anxiety among ZTTK patients (https://zttksonshinefoundation.org/). Additionally, 6 out of 46 individuals examined exhibited autistic spectrum disorders or behaviors, with this feature being more prominent in adults (50%) than younger individuals (under 18 years old) (11).

To investigate whether *Son^+/−^* mice show any phenotypic traits associated with cognitive and behavioral abnormalities, we conducted tests on a cohort of 8-10 month-old mice. We first performed the open field test (Figure 2E), which measures overall locomotor activity, exploratory habits, and anxiety-like behavior (27, 28). We tracked mouse activity across an “open field” arena with a center inner zone (Figure 2F). Compared to WT mice, *Son^+/−^* mice covered a significantly greater distance and had higher velocity (Figure 2, G and H), indicating hyperactivity or anxiety-related behaviors. We also measured thigmotaxis, which is the tendency to remain close to walls and is associated with anxiety-related behaviors (29). Interestingly, *Son^+/−^* mice spent more time close to the wall and less time in the open field compared to WT mice (Figure 2, F and I), suggesting that *Son^+/−^* mice have anxiety-like characteristics.

Next, we evaluated whether *Son^+/−^* mice exhibited signs of cognitive impairment using the Y-maze and novel object recognition tasks. In the Y-maze test, mice should be able to memorize the previously visited arm and prefer to explore new arms (a phenomenon known as “spontaneous alternation”) (30). We found that *Son^+/−^* mice exhibited a significant decrease in the percentage of spontaneous alternations (Figure 2J) and an increased frequency of repeatedly visiting the same arm (repeated arm errors) (Figure 2K), indicating deficits in spatial short-term working memory. To further assess their learning ability and memory, we employed the novel object recognition test (Figure 2L). Since mice have innate preference of novel object over a previously explored, familiar object, they typically spend more time at the novel object over the familiar object (31). Interestingly, while WT mice spent more time with the novel object than the previously explored object, *Son^+/−^* mice did not show this innate preference and spent less time exploring the novel object and more time with the familiar object (Figure 2, M and N), indicating an impairment in preferential recognition of the novel object.

Our behavior assessments suggest that our *Son^+/−^* mice could be a useful model for studying ZTTK syndrome-associated intellectual disability and behavior disorders, including anxiety-like behavior and cognitive/learning impairment, and for identifying how *Son* haploinsufficiency contributes to these behavior patterns and cognitive impairments.

### *Son^+/−^* mice recapitulate many other clinical features of human ZTTK syndrome patients, including scoliosis/kyphosis, kidney hypoplasia/agenesis, and mild microcephaly

Another common phenotype observed in ZTTK syndrome patients is spine curvature, such as scoliosis and kyphosis (6, 7, 11). Alizarin Red/Alcian Blue staining of skeletons from 1-day-old pups showed curved spines in *Son^+/−^* mice, which mimic scoliosis and kyphosis while no significant abnormalities in ossification was found (Figure 3A). Micro-computed tomography (µCT) imaging of 8-week-old *Son^+/−^* mice showed spine curvature, which became more severe as they aged (26 weeks), particularly for the cervical and thoracic curves (Figure 3B).

**Figure 3.**
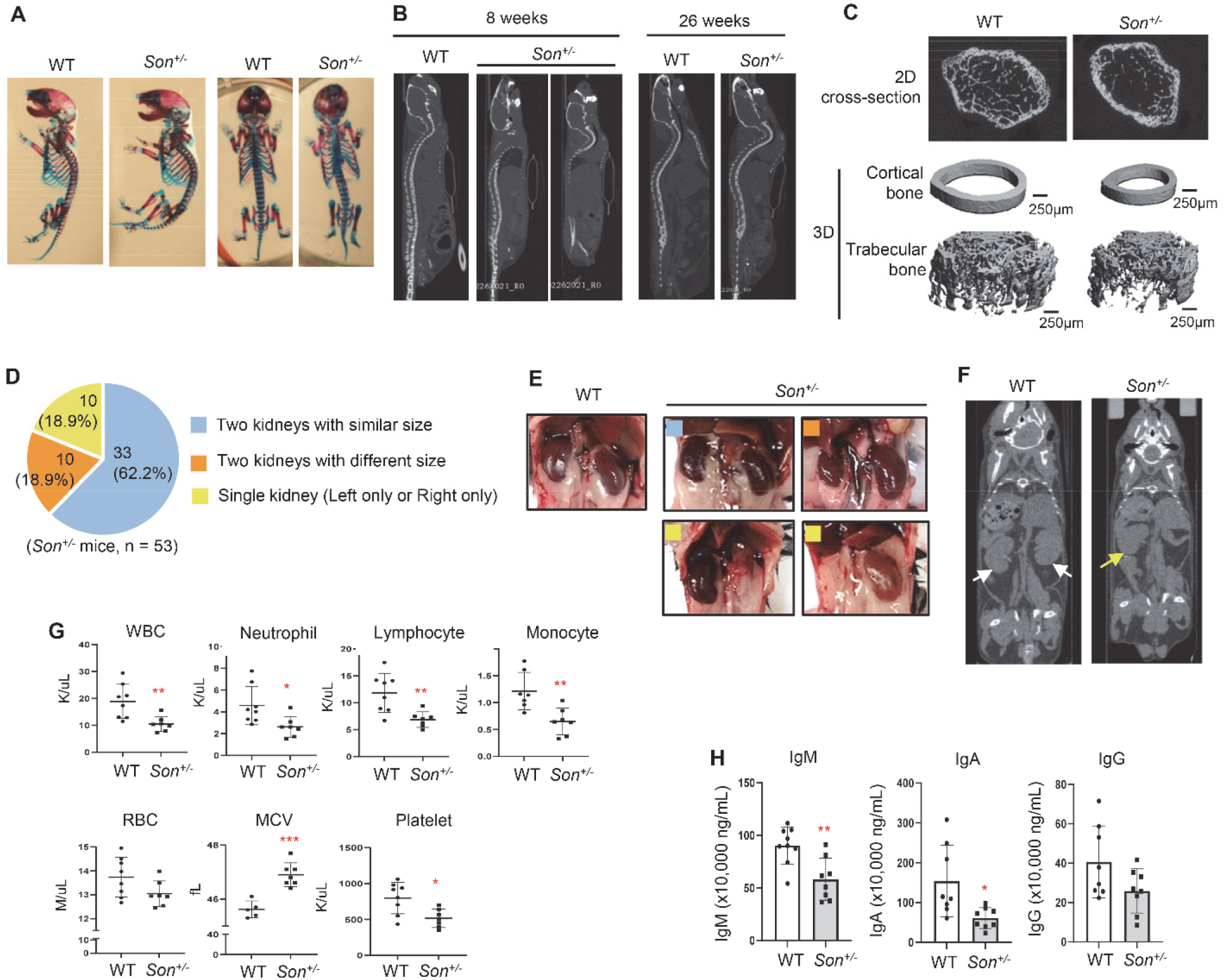
*Son^+/−^* mice faithfully recapitulate clinical features observed in human ZTTK syndrome patients, including scoliosis/kyphosis, kidney hypoplasia/agenesis, and hematological abnormalities. (**A**) Alizarin Red and Alcian Blue staining of WT and *Son^+/−^* mouse skeletons (postnatal day 1) showing signs of scoliosis and kyphosis. (**B**) Sagittal view of the µCT images showing kyphosis in *Son^+/−^* mice, which worsened as they aged. (**C**) 2D section and 3D reconstruction for the µCT scan of the femurs of 6 week-old WT and *Son^+/−^* mice. (**D**) Pie chart indicating the percent of the indicated kidney status determined in total 53 *Son^+/−^* mice. (**E**) Representative images of normal kidneys from WT mice, and normal and abnormal kidneys observed in *Son^+/−^* mice. Color-coded squares shown (**D**) were marked on *Son^+/−^* kidney images to indicate the kidney types. (**F**) Coronal view of the µCT images of WT and *Son^+/−^* mice, indicating presence of two kidneys in a WT mouse (white arrows) and one kidney in a *Son^+/−^* mouse (a yellow arrow). (**G**) Complete blood counts (CBC) from whole blood of WT and *Son^+/−^* mice at 13 weeks of age. (**H**) ELISA analysis of immunoglobulins (IgM, IgA, and IgG) in the plasma of WT and *Son^+/−^* mice. For (**G)** and (**H**), data are presented as mean ±SD, n=7-8. *p < 0.05, **p < 0.01, ***p < 0.001.

To examine the bone structure, the distal femoral metaphysis was examined by 2D and 3D reconstruction from µCT images (Figure 3C). The results revealed a significant reduction in the trabecular bone volume fraction (BV/TV) in *Son^+/−^* mice when compared to control littermates (Supplemental Figure 4). A decrease in trabecular number (Tb.N) and an increase in trabecular spacing (Tb.Sp) were also observed in *Son^+/−^* mice, particularly in the TRI plate model of analysis (Supplemental Figure 4). Based on similar trabecular thickness (Tb.Th) between groups, this indicates that reduced Tb.N resulted in the reduced trabecular bone volume fraction in *Son^+/−^* mice. During cortical bone analysis in the mid-shaft of the femur of *Son^+/−^* mice, the µCT images also show that the cross-sectional area of the bone was smaller than that of WT mice (Figure 3C). These findings revealed that *Son^+/-^* mice have reduced bone mass in addition to spinal curvature.

Previously, our group reported kidney abnormalities found in human ZTTK syndrome patients, including horseshoe kidneys, single kidney, and early onset of kidney cysts (10). Among the 53 *Son^+/−^* mice we analyzed for kidney features, 33 out of 53 mice had two kidneys of similar size, 10 mice (18.9%) had a single kidney (left or right only), and 10 mice (18.9%) had two kidneys with one hypoplastic kidney (Figure 3, D, E and F). We did not observe any mice with horseshoe kidneys or kidney cysts, and H&E staining showed minimal to mild tubular generation/necrosis and mild interstitial edema in the samples from mice with a single kidney (Supplemental Figure 5). Our previous analysis indicated that 37.38% of ZTTK syndrome patients previously examined for their kidneys (17 out of 45 patients) showed abnormal renal morphology (11). Interestingly, 37.73% of the *Son^+/−^* mice we examined (20 out of 53) had a single kidney or one hypoplastic kidney (Figure 3F), indicating a very similar level of phenotype penetrance in humans and mice.

Furthermore, MRI of the brain showed that *Son^+/−^* mice have decreased total brain volume compared to WT littermates (Supplemental Figure 6), which recapitulates the microcephaly phenotype observed in 11 out of 40 (25%) ZTTK syndrome patients reporting their head circumference (11).

Taken together, these findings indicate that *Son^+/−^* mice are an animal model that can recapitulate the skeletal, renal, and brain phenotypes observed in human ZTTK syndrome patients and could be valuable research tools to study these abnormalities.

### A wide spectrum of hematological abnormalities has been observed in children with ZTTK syndrome, and *Son^+/−^*mice showed similar abnormalities in peripheral blood analysis and immunoglobulin measurements

Among the clinical symptoms observed in individuals with ZTTK syndrome, a clinically important but not well-defined issue is their hematological/immunological symptoms. Immunoglobulin deficiency and frequent infections were previously reported by our group and others (6, 7, 11), and 15 out of 46 published cases (32.6%) indicated abnormalities in the immunological and/or hematological system, such as immunoglobulin deficiency, particularly IgA and IgG, and frequent and severe infections (11). Voluntarily reported clinical features from families to the ZTTK SON-Shine Foundation include not only immunoglobulin deficiency, poor responses to vaccines, recurrent infections, and hyper-reactivity to the common cold, but also a wide range of other hematological abnormalities, including bone marrow failure, high mean corpuscular volume (MCV), polycythemia, severe anemia, and thrombosis. Some children had to receive a transfusion due to low platelet levels. Additionally, low WBC counts, especially low levels of lymphocytes, neutrophils, and monocytes, have been identified from the complete blood count (CBC) (https://zttksonshinefoundation.org/; Summarized in Supplemental Table 1). Interestingly, CBC analyses of *Son^+/−^* mice identified significant decreases in WBC, and low levels of lymphocytes, neutrophils, and monocytes. RBC and hemoglobin levels were normal, but high MCV levels and low platelet counts in *Son^+/−^* mice were observed (Figure 3G and Supplemental Figure 7). Another noticeable hematopoietic feature reported from ZTTK syndrome patients is low levels of immunoglobulins, specifically IgM, IgA, and IgG, (7, 11) (also documented in https://zttksonshinefoundation.org), and we indeed observed significant decreases in these immunoglobulins, especially IgM and IgA, in *Son^+/−^* mice even in the steady state (Figure 3H). These findings indicate *Son^+/−^* mice recapitulate the CBC abnormalities found in human ZTTK syndrome patients.

### *Son^+/−^* fetal liver hematopoiesis shows phenotypic alterations of hematopoietic stem and progenitor cells with a decreased pool of early stage HSPCs and an increased bias toward megakaryocyte/erythroid lineages

To understand that the hematopoietic features found in ZTTK syndrome patients are due to *SON* LoF, we further analyzed hematopoiesis of our *Son^+/−^* mice. Since many hematological abnormalities can be attributed to altered phenotypes of early stage hematopoietic stem and progenitors (HSPCs), we first analyzed HSPCs during fetal liver hematopoiesis. While fetal liver cellularity does not show significant changes (Figure 4A), we found that the percentage of early stage HSPCs (defined by absence of lineage markers and positivity of cKit and Sca1 expression; Lin*^−^*, Sca1^+^, cKit^+^ cells; denoted as LSK cells) was significantly decreased in the *Son^+/−^* fetal liver (Figure 4, B, C, and D, and Supplemental Figure 8A).

**Figure 4.**
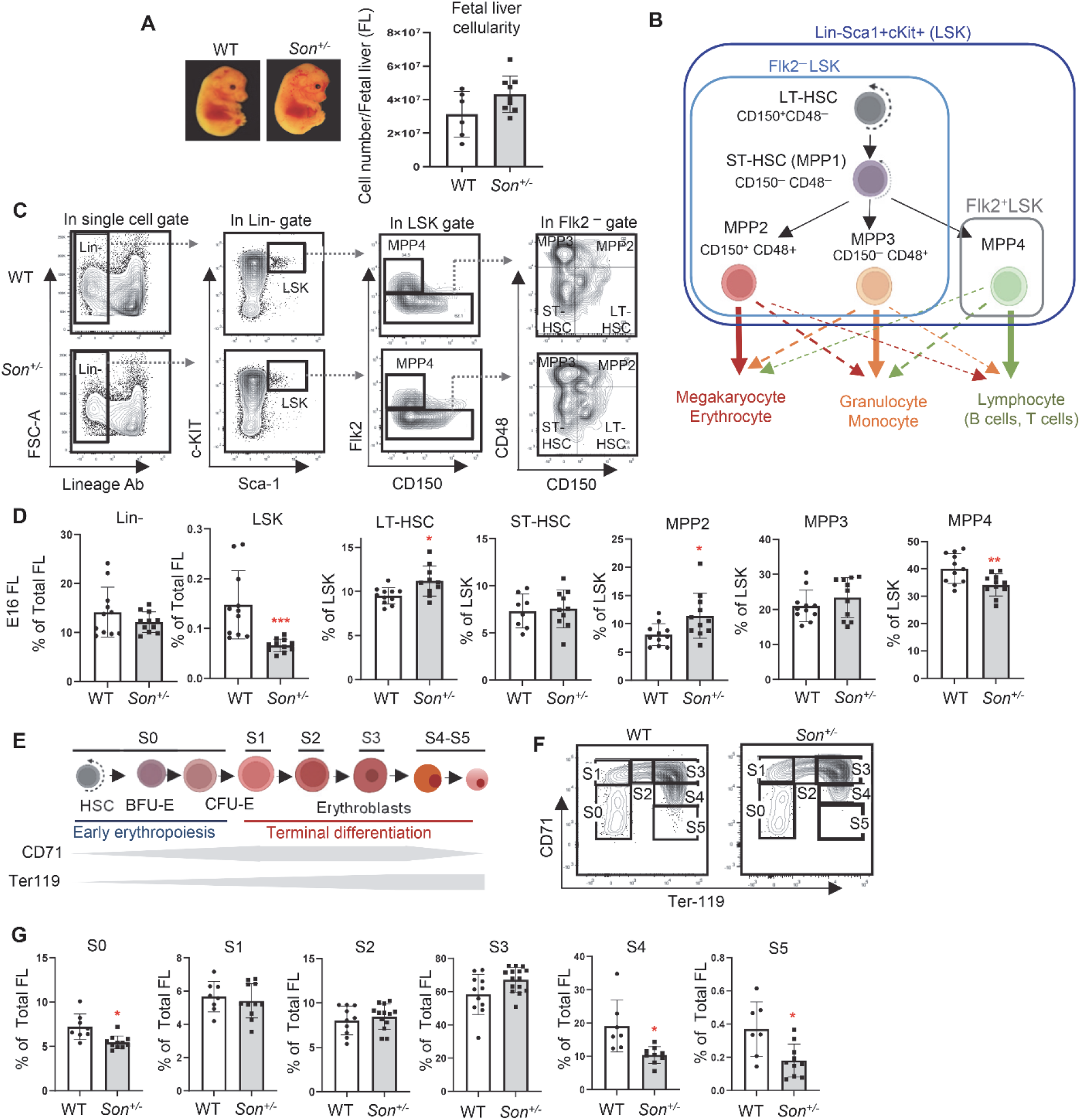
*Son* haploinsufficiency causes a reduction of the early stage HSPC (LSK) pool size, an imbalance of myeloid vs. lymphoid lineage-biased MPPs, and impaired erythroid terminal differentiation during fetal liver hematopoiesis in *Son^+/−^* embryos. (**A**) Representative image of WT and *Son^+/−^* embryos at E14 (left panel). Fetal liver cellularity at E14 (right panel) showing no statistically significant difference between WT and *Son^+/−^* fetal livers. (**B**) A schematic indicating subpopulations within the LSK population (LT-HSC, ST-HSC, MPP2, MPP3, and MPP4), their lineage bias, and the surface markers. (**C**) Representative flow cytometry contour plots showing the gating scheme used to identify LSK and other subpopulations in the fetal liver. (**D**) Frequency of the indicated populations within total fetal liver cells (for Lin- and LSK) or within the LSK population (LT-HSC, ST-HSC, MPP2, MPP3 and MPP4). (**E**) A schematic depicting the process of erythroid differentiation and the expression levels of CD71 and Ter119. (**F**) Representative flow cytometry contour plots demonstrating fetal liver erythroid subsets S0-S5 indicated by CD71 and Ter119 expression. (**G**) Frequency of the indicated erythroid subsets, presented as percent of total fetal liver cells. For (**A**), (**D**) and (**G**), data are presented as mean ± SD, n=8-11, *p < 0.05, **p < 0.01, ***p < 0.001.

Next, to further characterize the HSPC population, phenotypic hematopoietic stem cells (HSCs) and lineage-biased multipotent progenitors (MPPs) were analyzed from the *Son^+/−^* fetal liver using Flk2 and two SLAM (signaling lymphocyte activation molecule) family markers, CD48 and CD150 (32) (Figure 4, B and C). We observed a mild increase in long-term hematopoietic stem cells (Flk2*^−^* CD150^+^ CD48*^−^* LSK; denoted as LT-HSC) in the *Son^+/−^* fetal livers. More interestingly, among lineage-biased MPPs, megakaryocyte/erythroid (MegE) lineage-biased MPPs (Flk2^−^ CD150^+^ CD48^+^ LSK; denoted as MPP2) were increased, while lymphoid lineage-biased MPPs (Flk2^+^ CD150^−^ LSK; denoted as MPP4) were decreased in both E14 and E16 *Son^+/−^* fetal livers (Figure 4, C and D, and Supplemental Figure 8A). These changes were marginal, but statistically significant, suggesting that HSPC subpopulations were already altered during fetal liver hematopoiesis.

Knowing that *Son^+/−^* fetal liver HSPCs contain an increased level of MegE-biased MPPs, we further analyzed the downstream myeloid progenitors (Lin*^−^* cKit^+^ Sca1*^−^* ; denoted as LK (Sca1^-^) cells) based on a classical model of myeloid progenitor differentiation (33) (Supplemental Figure 8B). We found that the LK (Sca1^-^) population is decreased, and the megakaryocyte/erythroid progenitor (MEP) population showed an increase, (Supplemental Figure 8, C and D). Together with the MPP2 increase within the LSK population and the MEP increase within the LK population, our data suggest that *Son* haploinsufficiency leads to an early stage HSPC lineage disposition towards the MegE lineage during fetal liver hematopoiesis.

### Terminal erythroid differentiation is impaired during fetal liver hematopoiesis in *Son^+/−^* **embryos.**

One of the hematological features voluntarily reported to the ZTTK SON-Shine foundation is the increased size of erythrocytes (elevated value of mean corpuscular volume; MCV), and we also identified this feature from CBC of *Son^+/−^* mice (Figure 3H). During erythroid terminal differentiation, erythroblasts undergo maturational cell divisions in which cells become smaller. Therefore, enlarged erythrocytes could be indicative of impaired terminal differentiation. From the fetal liver, we examined erythroid lineage cells in 6 different stages (S0 – S5) based on CD71 (transferrin receptor) and Ter-119 expression (Figure 4E). Our analysis revealed that erythroid progenitors with colony-forming potential (BFU-E and CFU-E) and erythroblasts up to the S3 stage showed similar levels of cell numbers in the WT and *Son^+/−^* fetal liver. However, we found a significant reduction in S4 and S5 stages in the *Son^+/−^* fetal liver (Figures 4F and 4G).

These results indicate that early stage erythroid progenitors are intact, and onset of erythroid differentiation normally occurs in the *Son^+/−^* fetal liver. However, the late stage of terminal differentiation with downregulation of CD71 is impaired. Taken together, our analyses defined impaired erythroid terminal differentiation as a phenotype caused by *Son* haploinsufficiency.

### The bone marrow of *Son^+/−^* mice showed a reduction in early stage HSPC population and altered lineage bias with expansion of myeloid-lineage biased MPPs and reduction of lymphoid-biased MPPs

Next, we analyzed the bone marrow of *Son^+/−^* mice (8-10 weeks), which showed an overall reduction of bone marrow cellularity compared to that of WT mice (Figure 5A). We found that the portion of LSK cells (early stage HSPCs) was decreased in the bone marrow of *Son^+/−^* mice (Figure 5, B and C), which is similar to our observation from the fetal liver (Figure 4D). In addition, within the LSK population of *Son^+/−^* mice, we found an expansion of MegE-biased MPP2 and granulocyte/monocyte (GM)-biased MPP3 and a reduction of lymphoid lineage-based MPP4 (Figure 5C). These data, together with our fetal liver analyses, revealed that *Son* haploinsufficiency decreased the overall size of early stage HSPCs (LSK cells) and there is a skewed bias in the MPP lineage towards the myeloid lineages (MegE and GM lineages) rather than the lymphoid lineage. Particularly, MPP2 expansion and MPP4 reduction were already initiated during fetal liver hematopoiesis and persisted in the adult bone marrow hematopoiesis.

**Figure 5.**
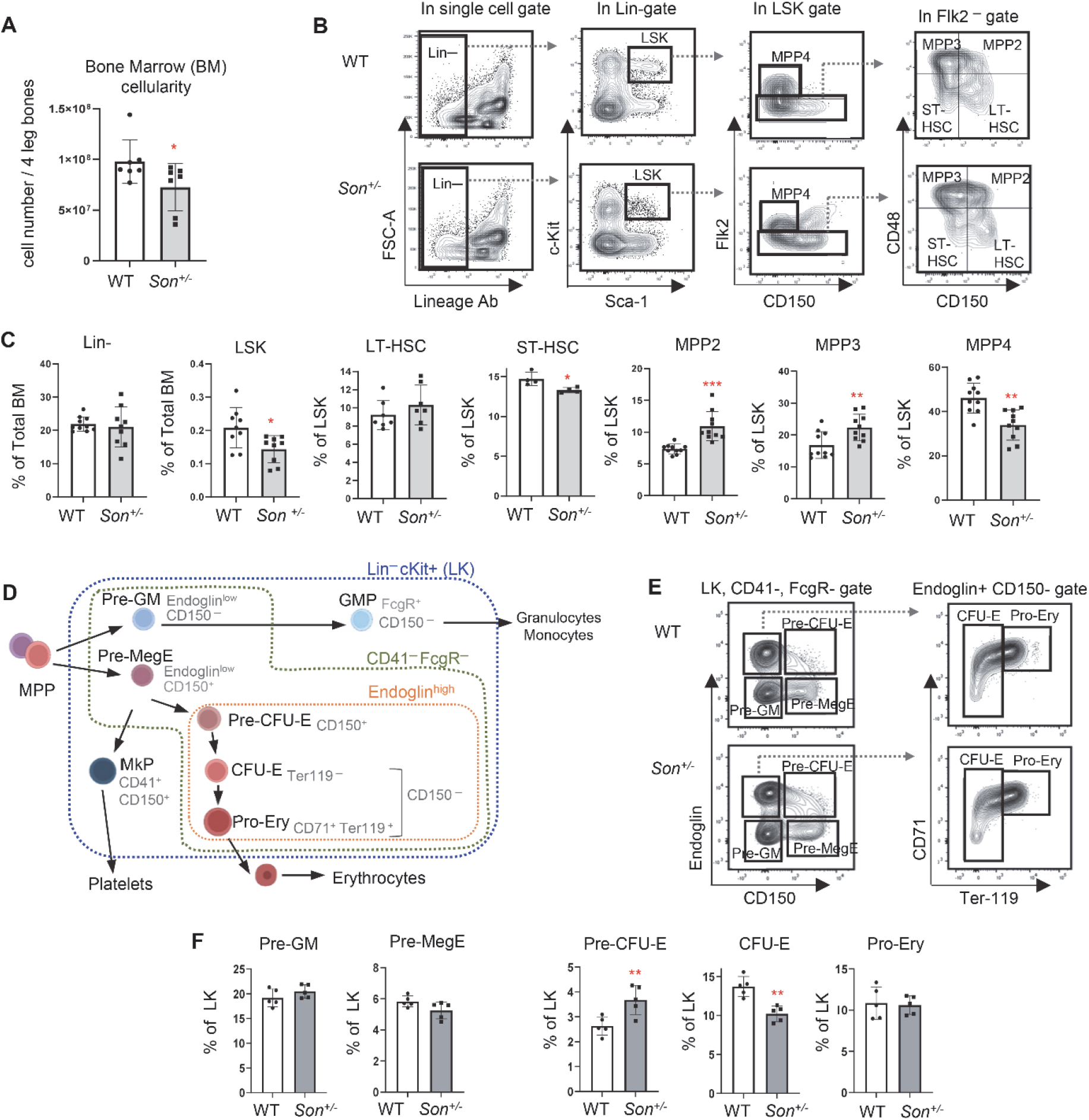
*Son* haploinsufficiency causes a reduction of the early stage HSPC (LSK) pool size, increased myeloid lineage-based MPPs, decreased lymphoid-biased MPPs, and increased early-stage erythroid progenitors in the bone marrow of adult *Son^+/−^* mice. (**A**) Bone marrow cellularity from 8–10-week-old mice, presented as mean ±SD. n=7, *p < 0.05. (**B**) Representative flow cytometry contour plots showing the gating scheme used to identify LSK (Lin-Sca1+ cKit+) and other subpopulations within the LSK pool of adult bone marrow. (**C**) Frequency of bone marrow LSK subsets presented as mean ± SD, n=9, *p < 0.05, **p<0.01, ***p<0.001. (**D**) A schematic depicting differentiation of myeloerythroid progenitors (Lin-cKit+ [LK] cells) and surface markers defining the lineages and stages. (**E**) Flow cytometry contour plots demonstrating the gating scheme to define myeloerythroid progenitor subpopulations within CD41-FcgR-LK cells, using Endoglin, CD150, CD71, and Ter119 markers. (**F**) Frequency of bone marrow myeloerythroid subsets within LK cells, presented as mean ± SD, n=5, *p < 0.05, **p < 0.01.

Knowing that myeloid lineage-biased MPPs are increased in *Son^+/−^* bone marrow hematopoiesis, we further examined myeloid progenitor development. Using CD150, FcgR, endoglin (CD105), CD41, CD71, and Ter119, we delineated the hierarchy of functionally distinct myelo-erythroid progenitor cells within Lin^─^ Sca1^─^ cKit^+^ (LK) cells (34) (Figure 5D). We did not see notable changes in granulocyte/monocyte precursors (pre-GM and GMP) and megakaryocyte progenitors (MkP). However, we found that early stage erythroid progenitors with colony forming potential (pre-CFU-E) is increased while the next stage progenitors, CFU-E, are decreased in the *Son^+/−^* bone marrow (Figure 5, E and F, and Supplemental Figure 9, A and B). Taken together, these analyses revealed that while *Son^+/−^* mice have an increase in MegE lineage-biased MPPs (MPP2), they have a compromised ability to move through the erythroid-lineage differentiation process.

### *Son^+/−^* mice show impaired B cell maturation in the spleen

One of most prominent features of hematological issues of ZTTK syndrome patients is their low immunoglobulin levels, which was recapitulated by our *Son^+/−^* mice (Figure 3H). To address whether this is attributed to impaired B cell development in *Son^+/−^* mice, we first analyzed series of B cell subsets in the bone marrow (Hardy fractions) (35) as well as spleen (Figure 6A and Supplemental Figure 9, C and D). While bone marrow B cells are mostly intact and not altered in *Son^+/−^* mice, IgD expression was slightly reduced within the CD43^+^ late state developing B cell population, resulting in increased Small Pre-B fraction (fraction D) and decreased recirculating B fraction (fraction F) (Figure 6B and Supplemental Figure 9, C and D). The most interesting alteration was observed in CD93 expression analysis. CD93 (AA4.1) is expressed in immature or “transitional” B cells in the spleen, but it is low or devoid in mature B cells (36) (Figure 6C). We found that CD93^+^ transitional B cells are significantly increased, and in contrast, CD93*^−^* naïve mature B cells are decreased in the *Son^+/−^* mouse spleen (Figure 6, D and E).

**Figure 6.**
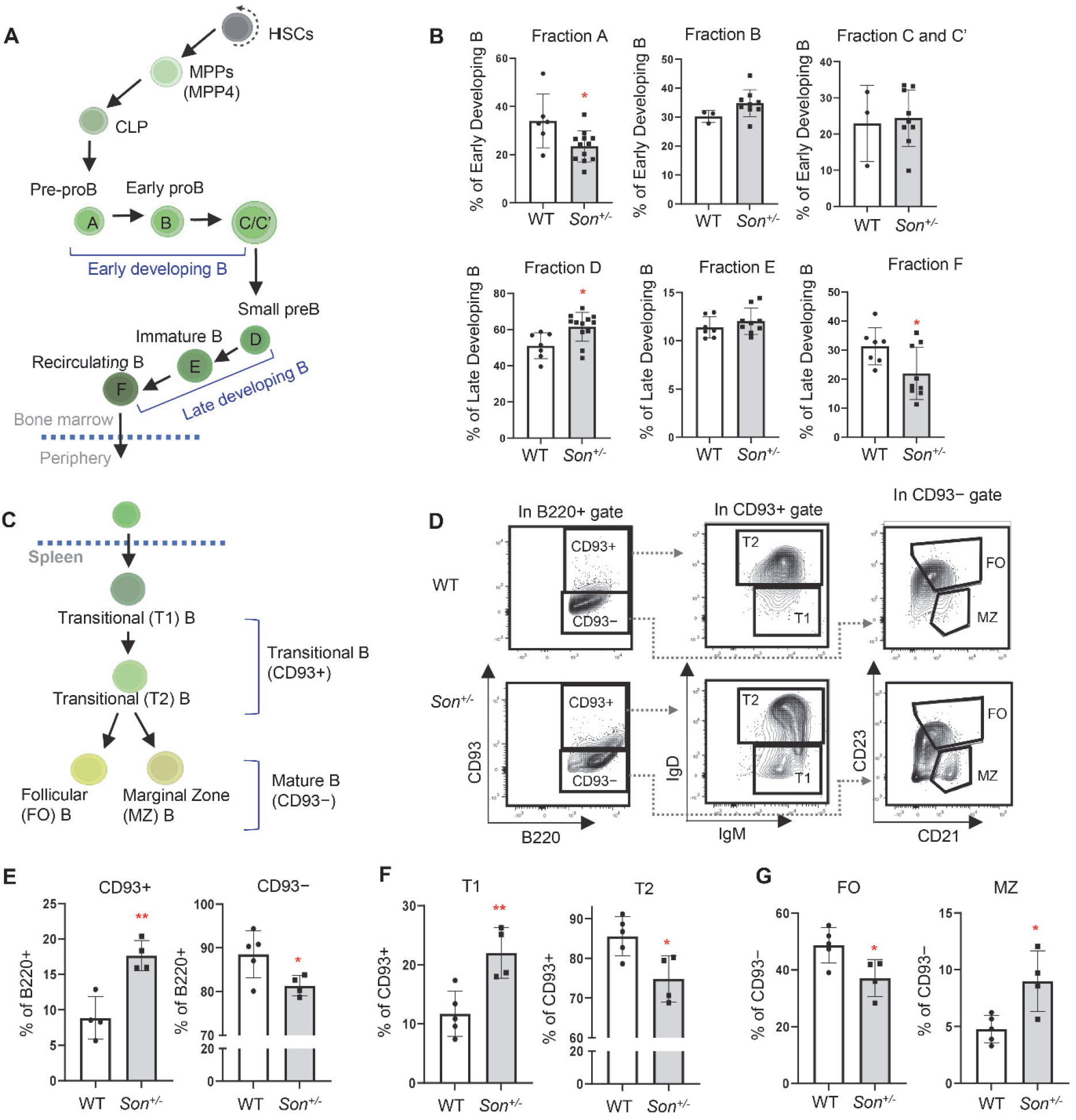
*Son* haploinsufficiency causes impaired B cell differentiation in the spleen of *Son^+/−^* mice. (**A**) A schematic depicting B cell development in the bone marrow. (**B**) Frequency of bone marrow B cell subsets (Hardy fractions) determined by flow cytometry analysis of surface markers. Data are expressed as mean ± SD, n=3-12, *p < 0.05, **p < 0.01. (**C**) A schematic depicting B cell differentiation into transitional B and subsequently mature naïve B cells in the spleen. (**D**) Flow cytometry contour plots demonstrating the gating scheme of CD93^+^ (transitional) and CD93^−^ (naïve mature) B cells within the spleen B220^+^ cell population, T1 and T2 transitional B cells within the CD93^+^ population, and FO and MZ B cells within the CD93^−^ population. (**E, F** and **G**) Frequency of the indicated spleen B cell subsets, expressed as mean ±SD, n=4-5, *p < 0.05, **p<0.01.

Within transitional B cells, transitional T1 cells differentiate into transitional T2 cells, which are the immediate precursor of the mature B cells (37). We further analyzed T1 and T2 cells using the IgM and IgD expression levels, and found that IgD^low^ T1 cells are increased while IgD^high^ T2 cells were decreased (Figure 6, D and F). These results indicate that within transitional B cells, the developing process from the “T1” subset to the “T2” subset is hindered, resulting in a blockage that prevents transitional B cells from progressing to naïve mature B cells. Further analyses of the CD93^−^ naïve B cell population demonstrated a reduced fraction of follicular (FO) B cells (CD23^+^ CD21^int^ IgM^low^ IgD^high^) and an increased marginal zone (MZ) B cell fraction (CD23^−^ CD21^high^ IgM^high^ IgD^low^) in *Son^+/−^* mice (Figure 6, D and G). These findings revealed significant perturbations in the normal early B cell maturation processes within the spleen of *Son^+/−^* mice, which are likely contributing factors to the observed low levels of immunoglobulins in both *Son^+/−^* mice and human ZTTK syndrome patients.

### Single-cell profiling and transcriptional state-based clusters of hematopoietic stem and progenitor cells reveals significant decreases of lymphoid/B cell lineage clusters in *Son^+/−^*mice

Our flow cytometric analysis of cell surface marker expression strongly suggests that *Son^+/−^* mice exhibit significant phenotypic changes in their hematopoietic stem and progenitor cells. In order to obtain a more comprehensive understanding of these changes at the molecular level, we employed single-cell RNA-sequencing (scRNAseq) using both WT and *Son^+/−^* bone marrow HSPCs isolated and sorted based on their Lin*^−^* cKit+ (LK) cell markers (24,355 total LK cells from 2 WT mice; 31,232 total LK cells from 2 *Son^+/−^* mice; Supplemental Figure 10, A and B). By conducting an in-depth analysis of the transcriptional states (Appendix 1) and cell cycle phases of individual cells (Supplemental Figure 10C), we identified 22 distinct clusters that represent the various hematopoietic stem and progenitor populations and their lineage trajectories (Figure 7A).

**Figure 7.**
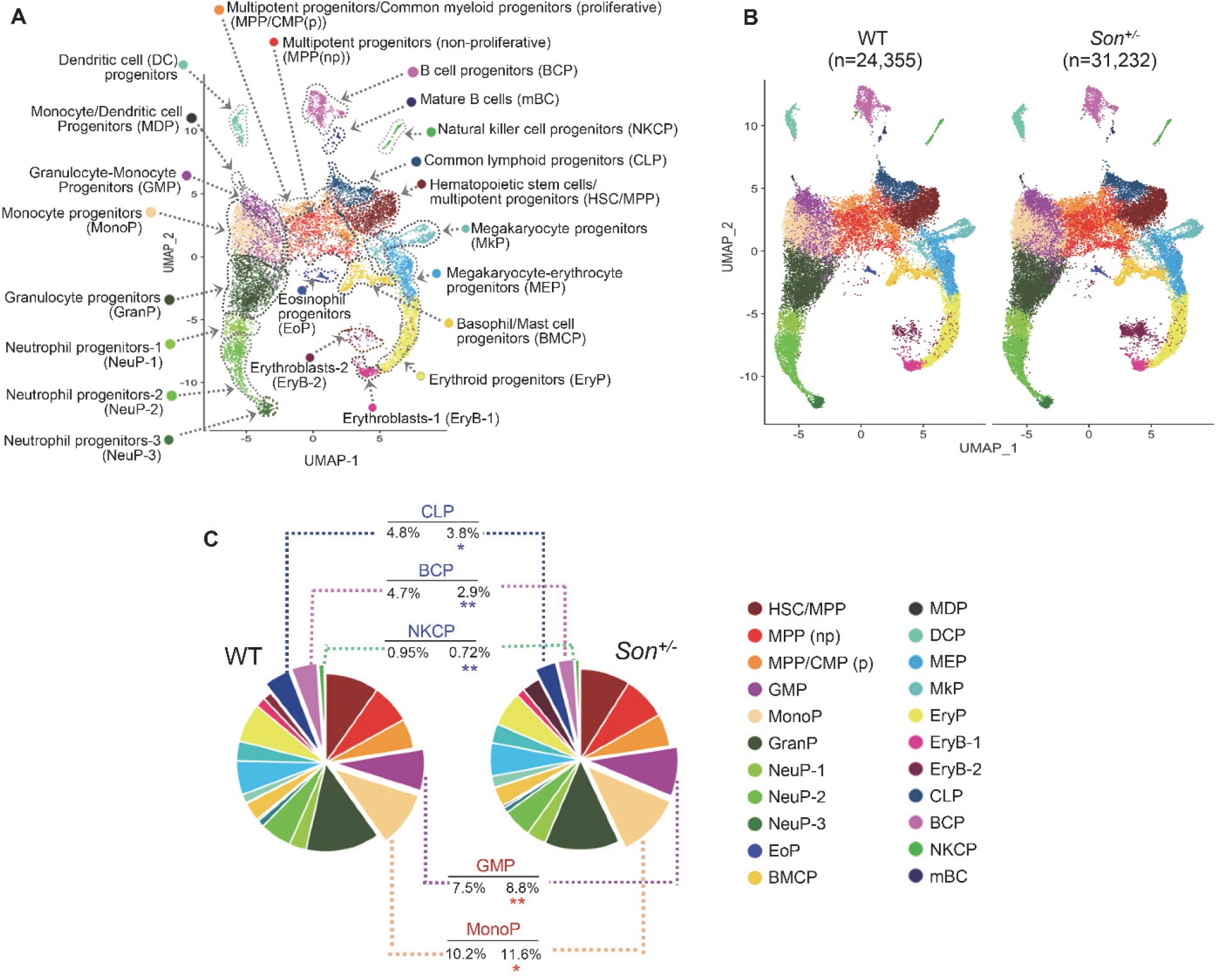
Single-cell RNA-sequencing and transcriptional state-based clustering of hematopoietic stem and progenitor cells reveals significant decreases of lymphoid lineage clusters in *Son^+/−^* mice. (**A**) UMAP plot showing 22 cell clusters identified from scRNAseq of mouse bone marrow Lin-c-kit+ (LK) cells. Cell clusters are annotated based on the enriched genes and cell cycle status. (**B**) UMAP plots showing the 22 clusters, as described in (**A**), identified in the bone marrow LK cells from WT and *Son^+/−^* mice. (**C**) Pie-chart illustrating the color-coded cell populations of the percentages of each cell cluster within total annotated cells. Clusters that showed either significantly increased (red asterisk) or decreased (blue asterisk) in *Son^+/−^* compared to WT are indicated. *p < 0.05, **p < 0.01.

While the HSPCs of both WT and *Son^+/−^* mice share similar cluster maps (Figure 7B), we observed several notable differences in the sizes of certain clusters within LK cells. We found that *Son^+/−^* mice have an increased proportion of GMP and monocyte progenitors (MonoP) (Figure 7C), and, of particular interest, we observed significant reductions in the size of clusters containing lymphoid lineage cells in *Son^+/−^* mice. These clusters include common lymphoid progenitor (CLP), B-cell progenitors (BCP), and natural killer cell progenitors (NKCP) and (Figure 7C). Among these, B cell progenitors exhibited the most noticeable reduction (38% reduction; 4.67% of WT annotated LK cells vs. 2.90% of *Son^+/−^* annotated LK cells). These transcriptome-based clusters revealed that, consistent with our data from surface marker-based analysis, *Son* haploinsufficiency leads to a significant reduction in lymphoid lineage-committed progenitor cells and an increase in early stage myeloid lineage progenitors.

### Altered gene expression patterns were identified in several HSPC clusters in *Son^+/−^* mice, including decreased expression of lymphoid lineage/B cell development genes and aberrant upregulation of erythroid lineage-promoting genes

To investigate the impact of *Son* haploinsufficiency within specific hematopoietic stem and progenitor cell (HSPC) clusters, we conducted an analysis of differentially expressed genes (DEGs) in each cluster. Notably, we observed a significant downregulation of critical genes related to lymphoid lineage and B-cell lineage development in the CLP and BCP clusters of *Son^+/−^* mice compared to those in wild-type mice (Figure 8, A and B). These genes include *Il7r* (essential for lymphoid lineage cell development), *Ebf1* (a key transcription factor for B cell lineage commitment), *Ly6d* (required for early-stage B cell specification), *Rag1* (critical for immunoglobulin gene rearrangement), and *Cd79a* (a critical component of B cell receptor signaling) (38–43). For these genes, we observed not only decreased expression levels per cell (Figure 8A), but also the decreased percentage of the cells expressing the assigned gene within a cluster (Figure 8B). The downregulation of these genes may contribute to the reduction in CLP and BCP cluster size and suggest potential defects in the differentiation and function of B cell lineage cells.

**Figure 8.**
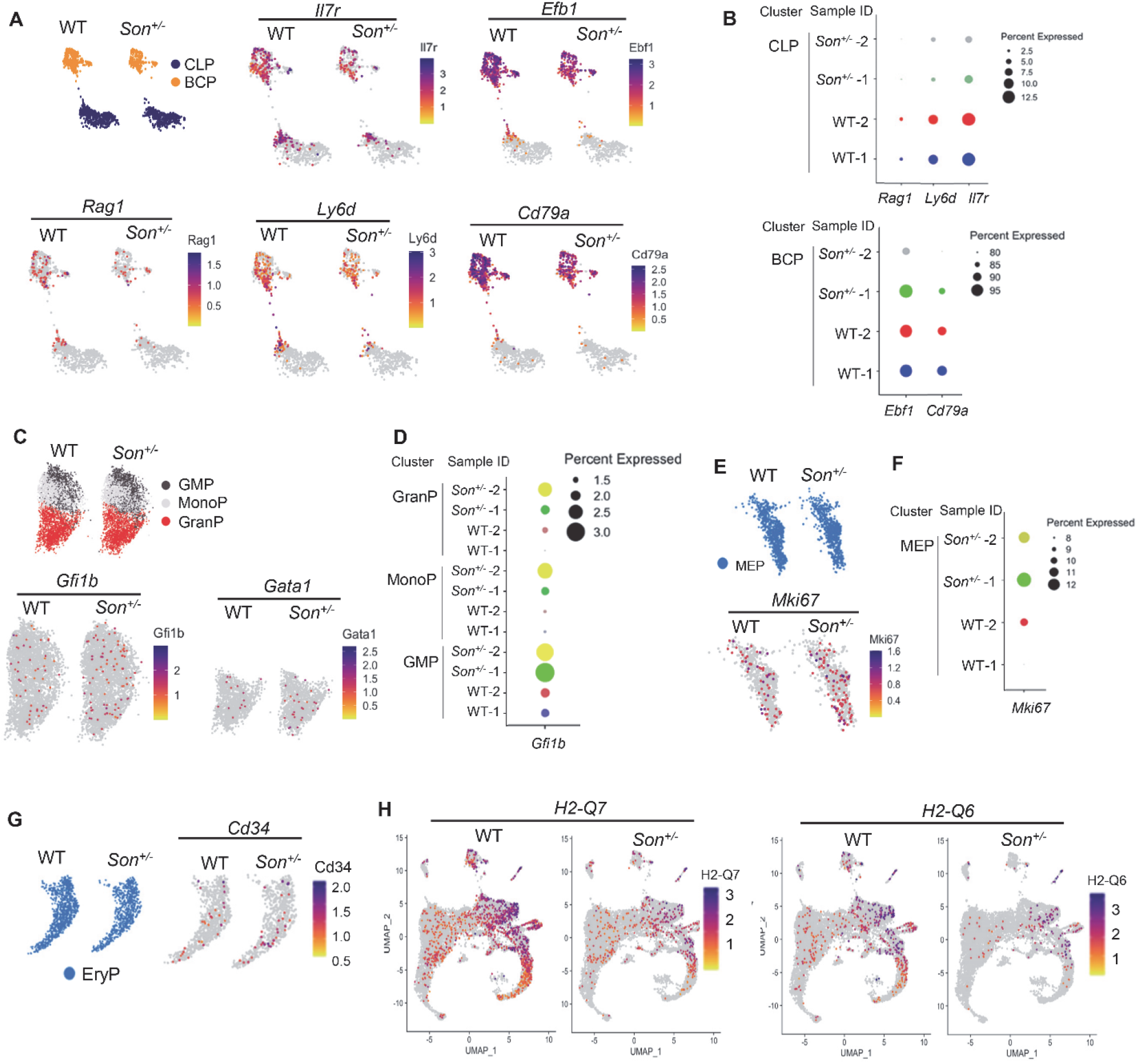
Differential gene expression analysis in hematopoietic progenitor clusters identified altered expression of genes critical for lymphoid/B cell lineage development and aberrant activation of myeloid lineage-regulating genes in *Son^+/−^*mice. (**A**) Feature plots showing the common lymphoid progenitor (CLP) and B cell progenitor (BCP) clusters from WT and *Son^+/−^* mice and the expression distribution of indicated genes critical for lymphoid lineage specification and B cell development. (**B**) Dot plots visualizing the percent of the cells expressing the indicated genes. The size of the dot encodes the percentage of cells within the cluster. (**C** and **D**) Feature plots (**C**) and dot plots (**D**) showing the granulocyte/monocyte progenitor (GMP), monocyte progenitor (MonoP), and granulocyte progenitor (GranP) from WT and *Son^+/−^* mice, and the expression of the erythroid-lineage transcription factors. (**E** and **F**) Feature plots (**E**) and dot plots (**F**) of megakaryocyte/erythroid progenitor (MEP) expressing *MKi67*. (**G**) Feature plots showing Cd34 expression in the erythroid progenitor (EryP) cluster. (**H**) Feature plots showing the expression levels of *H2-Q7* and *H2-Q6* genes in all clusters from WT and *Son^+/−^* mice. For **A**, **C**, **E**, **G**, and **H**, expression levels for each cell are color-coded as indicated in the heatmap legend.

Interestingly, we observed abnormal upregulation of critical erythroid lineage transcription factor genes, such as *Gfi1b* (44) and *Gata1* (45), in the granulocyte and/or monocyte progenitor populations of *Son^+/−^* mice (Figure 8, C and D). This finding suggests that despite the increased cluster size of GMP and MonoP in *Son^+/−^* mice (Figure 7C), they exhibit increased expression of inappropriate lineage genes. In the MEP cluster, we detected significant upregulation of the *Mki67* gene (Figure 8, E and F), which encodes Ki-67, a marker of proliferation and active cell cycle (46). Since the acceleration of the cell cycle in MEP has been linked to fate specification toward the erythroid lineage (47), it is likely that the highly proliferative MEP in *Son^+/−^* mice is associated with an increased number of pre-CFU-E, as demonstrated in our phenotypic analysis (Figure 7, E and F). Additionally, we observed upregulation of *Cd34*, which is a maker enriched in immature progenitors (48), in the *Son^+/−^* erythroid progenitor cluster (EryP) (Figure 8G), suggesting impaired or reduced maturation occurring in the EryP of *Son^+/−^* mice.

In addition to the changes in lineage-specific gene expression, our DEG analysis revealed a significant downregulation of *H2-Q6* and *H2-Q7* in multiple cell types of *Son^+/−^* HSPCs, including the most upstream population in the hematopoietic hierarchy, HSC/MPP (Figure 8H). The *H2-Q6* and *H2-Q7* genes encode non-classical MHC molecules, which play a crucial role in antigen presentation and regulation of immune responses (49, 50).

Taken together, our analysis, based on transcription state-based clustering and gene expression, highlights the molecular mechanisms underlying decreased lymphoid-lineage progenitors, defective B cell development, increased early-stage myeloid progenitors with erythroid lineage potential, and impaired terminal differentiation. In addition to genes influencing hematopoietic lineage fate and differentiation, the downregulation of MHC molecules associated with innate immunity across multiple cell types could contribute to the observed weakened immune function in individuals with ZTTK syndrome.

## Discussion

Given recent discoveries on the pathogenic effects of compromised SON function in ZTTK syndrome, as well as the diverse cellular functions of SON in RNA splicing, transcription, and nuclear speckle organization, there is a pressing need for appropriate model organisms with disrupted SON expression. In this study, we created a genetically engineered mouse line of *Son* deletion and demonstrated that *Son* is indispensable for embryo development. Using the *Son^+/−^* mouse model, we identified the importance of the full gene dosage of *Son* in various organ development and hematopoiesis, which explains multiple phenotypes found in human ZTTK syndrome. Furthermore, our scRNAseq data provides a comprehensive view of the hematopoietic stem and progenitor landscape in *Son^+/−^* mice, and highlights the potential molecular mechanisms underlying the observed phenotypic alterations.

We emphasize the value of our mouse model in both biological and clinical aspects. First, our mouse model is a useful tool for the biological research to study SON’s role in organ development and cell-type specific gene expression. Partially due to its large gene size (∼7.3 kb for the coding sequence) and protein size (consisting with 2,426 amino acids), cellular functions of SON have not been under extensive research. In addition, the sequence analysis predicted that SON contains an intrinsically disorganized region (IDR) lacking any fixed structure (23), which poses challenges for structure-based functional studies. While our group has reported that SON mediates efficient RNA splicing and represses MLL-complex-associated transcriptional initiation, there could be many unknown biological functions of SON remaining unexplored. Recent studies have demonstrated that SON, together with SRRM2, forms a core of nuclear speckles to regulate the nuclear speckle integrity (23) and is involved in boosting p53-mediated gene expression (24). Interestingly, SON has been shown to be a critical factor regulating 12-hour rhythm of nuclear speckle phase separation dynamics (51). Considering these multiple roles of SON in cellular functions, a mouse model of *Son* knockout and haploinsufficiency is needed. In this study, we present the newly created mouse lines, *Son^flox/flox^* and *Son^+/−^*, which will serve as invaluable resources to study SON function in various organs as well as at different developmental stages. We show that homozygous knockout of *Son* (*Son^−/−^*) caused embryonic lethality, defining indispensable roles of *Son* in embryo development. In addition, using the *Son^+/−^* mouse model and the SON antibody we generated, we present the Son protein expression pattern in the whole body of developing embryos and several adult tissues and confirmed the specificity of our findings by comparing *Son^+/−^* embryos/mice with wild-type controls. These analyses present many outstanding questions regarding SON’s roles in different tissues/organs as well as the nature of diverse forms of SON proteins that we detected in Western blots.

The other aspect and more exciting point we emphasize here is the value and usefulness of the *Son^+/−^* mouse model for clinical research on human ZTTK syndrome. The pathogenic effect of *SON* LoF variants on causing this complex human genetic disease was first reported in 2016 by our group and others. These findings made major advances in clinical diagnosis and led to incredible family effort and participation, which launched an official patient/family-initiated foundation, the ZTTK SON-Shine Foundation. However, the roadblock that clinical practice for rare diseases is facing is the unclear definition of clinical symptoms due to low prevalence and lack of attention from research communities. This causes extreme frustration to families and difficulties for clinicians to guide the patients and families for follow-up cares. Our *Son^+/−^* mouse model recapitulates many of the clinical features of human ZTTK syndrome patients. Although more extensive studies will be required to fully delineate underlying molecular mechanisms, we present that our mouse model could serve as a useful tool to study each of these clinically significant features and develop therapeutics to improve or alleviate clinical symptoms.

Besides characterizing clinical features mentioned above, we highlighted hematopoietic features of *Son^+/−^* mice, which are closely linked to the symptoms found in ZTTK syndrome patients, and identified the altered hematopoietic lineage differentiation in *Son^+/−^* mice. We identified several critical changes in hematopoiesis caused by *Son* haploinsufficiency, including increased myeloid lineage-biased MPPs, especially megakaryocyte/erythroid lineage-biased MPPs (MPP2), and, in contrast, decreased lymphoid lineage-biased MPPs (MPP4) in early stage hematopoiesis. This finding is particularly interesting because expansion of myeloid lineage and reduction of lymphoid lineage early hematopoietic progenitor cells are typical features found in aged hematopoietic stem cells and are associated with various hematopoietic diseases, including myelodysplastic syndrome and myeloproliferative neoplasm (MPN) (52–57). Our study provides a significant guideline for clinical practice to ZTTK syndrome with hematopoietic involvement and suggests the need for monitoring myeloid expansion, including MDS and MPN, in these patients.

Interestingly, at the level of lineage-committed progenitor cells, we found an increase in the Pre-CFU-E population, which are erythroid lineage-committed precursors preceding CFU-E, in the phenotypic analysis of *Son^+/−^* mouse bone marrow. The increased Pre-CFU-E could be due to increased erythroid fate specification from the progenitors with megakaryocytic/erythroid lineage bi-potential (MEP or Pre-MegE) or a blockage of erythroid differentiation. Our scRNA-seq data, showing Mki67 upregulation in the MEP cluster and Cd34 upregulation in the EryP cluster, suggest that it is likely that both erythroid-favored lineage specification and impaired erythroid differentiation contribute to the Pre-CFU-E increase.

In our bone marrow LK cell scRNA-seq, the most significant impact of *Son* haploinsufficiency on hematopoietic gene expression was downregulation of genes critical for lymphoid development and B cell lineage development, such as *Il7r*, *Ebf1*, *Rag1*, *Ly6d* and *Cd79a*, in the transcriptionally defined CLP and BCP clusters. Together with surface marker-based phenotypic analysis showing impaired B cell maturation, our data suggest that the normal level of *Son* is required to express sufficient levels of genes directing lymphoid lineage specification and B cell development. Further investigations on terminal differentiation processes of each lineage and the origin of lineage bias will enable us to understand more about SON’s role in different hematopoietic lineages and how *SON* haploinsufficiency alters hematopoiesis in ZTTK syndrome patients. Our study supports the notion that CBC abnormalities and low levels of immunoglobulins should be considered as direct outcomes of SON LoF and these symptoms should be incorporated in the diagnosis process.

About 95% of rare diseases do not have treatments due to the lack of research on these diseases. Our attention to rare disease research is critical to improve the quality of patients’ lives. Our data presented here will elicit future research on ZTTK syndrome for individuals, particularly children, with this rare disease. We anticipate our study to be a key reference for the rare disease community and promote research on rare diseases.

## Methods

### Generation of *Son^flox/flox^* mice

*Son^flox/flox^* mice were generated by CRISPR/Cas9-based Extreme Genome Editing (EGE) system (Biocytogen). Briefly, two single guide RNAs (sgRNA) were designed to target the non-conserved intron regions flanking exon 2 (based on the Ensembl transcript ID ENSMUST00000114037.8; NCBI RefSeq NM_178880.4) within the mouse *Son* gene (NC_000082.6) located on mouse chromosome 16. The selected sgRNA and Cas9 mRNA were co-injected into C57BL/6 mouse zygotes together with the targeting vector containing homologous arms and the loxP element. The zygotes were then transplanted into pseudo-pregnant females to generate founder mice bearing the floxed *Son* allele. These mice were backcrossed to wild-type C57BL/6J mice (Jackson Laboratory; Strain # 000664) for at least 4 generations, followed by intercrossing to generate mice homozygous for the floxed allele (Son*^flox/flox^*).

### Generation of constitutive *Son^+/−^* mice

*Son^flox/flox^* mice were crossed to β-actin-cre (*ACTB-cre)* mice (Jackson Laboratory, Strain #019099; B6N.FVB-*Tmem163^Tg(ACTB-cre)2Mrt^*/CjDswJ) to obtain *ACTB-Cre^+^; Son^flox/wt^* mice. To obtain mice with heterozygous *Son* deletion in germline, *ACTB-Cre^+^; Son^flox/flox^* mice were backcrossed to wild-type C57BL/6J mice and offspring with excision of the floxed sequence within the *Son* gene in the absence of the *Cre* trans-gene were identified. The selected offspring were further backcrossed to wild-type C56BL/6J mice for more than 6 generations to ensure stable germline transmission of the *Son* allele with excision of deletion. The resulting mice have constitutive, heterozygous deletion of *Son* exon 2 from germline (*Son^+/−^*) without Cre expression, which causes a premature stop codon after encoding 9 amino acids in exon 3.

### Immunohistochemical staining of embryos

Formalin-fixed paraffin-embedded embryos section were heated (60 °C for 30 minutes) in oven, deparaffinized, and hydrated in xylene and 100%, 95%, 80%, 70% ethanol, and then in dH_2_O. After unmasking antigens in boiling citrate buffer (Sigma, C9999) with pH 6.0 for 40 minutes, the sections were incubated in 3% hydrogen peroxide for 10 minutes, rinsed with water, and blocked with 5% goat serum diluted in Tris-Buffered Saline (TBS) with 0.1% Tween-20 (TBST) for 30 minutes at room temperature (RT). Then, sections were incubated with the primary antibody, anti-SON generated against SON amino acids 77-84 (22), overnight at 4°C. After washing with TBST, sections were incubated in secondary antibody (anti-rabbit; Abcam AB205718) for 30 minutes at RT, and then with DAB (3,3’Diaminobenzidine) (Thermo PI34002) for 30 seconds. After incubation with Hematoxylin (Vector Laboratories, H3404) for 1 minute, slides were washed in dH_2_O, dipped in 0.3% ammonia water for 2 seconds and dehydrated in 70%, 80%, 95%, 100% ethanol and xylene. Images were taken using the Lionheart LX automated microscope (Agilent BioTek).

### Western blot analysis

Proteins from mouse tissue were extracted with 1xRIPA buffer (150 mM NaCl/1% sodium deoxycholate/1 mM EDTA/0.1% SDS/50 mM Tris pH 8/1% Triton X-100), Pierce Protease Inhibitor, EDTA-Free (Thermo Scientific, A32965), and Pierce Phosphatase inhibitor (Thermo Scientific, A32957). Samples were centrifuged at 13,000 xg for 30 mins at 4°C, and the supernatant protein samples were separated on 7% SDS-polyacrylamide gels and transferred onto PVDF membrane. Membranes were blocked in 5% BSA in 1x TBST (0.2%) for 1 hour at RT, and incubated with the respective primary antibodies overnight at 4°C. Antibodies that were used included anti-SON (22) and β-actin (Cell Signaling, 3700). Blots were washed in 1x TBST (0.2%) at RT and incubated with the secondary antibody for 2 hours on rotator at RT, following 3 times for 10 minutes/wash in 1x TBST (0.2%) at RT. Chemiluminescence detection was performed with the X-ray film (Alkali Scientific) and X-ray film processor (Konica).

### Alizarin red and alcian blue staining of skeletons

Neonatal mouse pups (day 1) were euthanized, and scalded in water (65°C) for 30 seconds and internal organs were removed, followed by fixing in 95% ethanol overnight and incubating in acetone overnight. The fixed samples were stained with Alcian blue overnight at RT and then destained by incubation in 95% ethanol overnight. Then, the samples were incubated in 1% KOH for 1 hour at RT and then incubated in Alizarin overnight at 4°C and then incubated in 50% glycerol: 50% (1%) KOH solution at RT until the specimen became clear. Skeletons were then transferred to 100% glycerol until images were taken.

### Microcomputed tomography (µCT)

Moue full-body mouse scans were performed under isoflurane in oxygen anesthesia and prospectively gated for single respiration phase using the following settings: 50kVp, 0.24 mA, 20ms exposure, 20 μm voxel size, using the U-CT^UHR^ µCT scanner (MiLabs, Utrecht, Netherlands). Scans were reconstructed using vendor software. For bone scans, femurs were isolated from 6-week-old female mice, skeletal muscle removed, and fixed in formalin for 48 hours followed by storage in 70% ethanol. Femurs were placed in a 12mm diameter scanning holder and imaged with the following settings: 70kVp, 114µA with an integration time of 200ms, and 500 projections per 180°, 12 µm voxel size, using the µCT40 desktop cone-beam scanner (Scanco Medical AG, Brüttisellen, Switzerland using µCT Tomography v6.4-2). Scans were reconstructed into 2-D slices and all slices analyzed using the µCT Evaluation Program (v6.5-2, Scanco Medical) and 3-D reconstruction performed using µCT Ray v4.2.

### Complete Blood Counts (CBC) and measurement of immunoglobulins

Peripheral blood counts were obtained by using heparin coated microvettes (Sarstedt), and heparinized blood was subjected to analysis using Hemavet 950FS for complete blood counts. IgM, IgG, and IgA titers in mouse plasma were measured using the ELISA kits from Bethyl Laboratories (IgM catalog #E99-101; IgG catalog #E99-131; IgA catalog #E99-103) according to the manufacturer’s instructions. The ELISA results were assessed with the BioTek SynergyHI microplate reader.

### Mouse behavior assessments

Open field test was conducted in a square Plexiglass box for 10 minutes, and the animal was tracked with an overhead camera. The automated tracking system (EthoVision) analyzed parameters such as distance moved, velocity, acceleration, and time spent in pre-defined zones. Y-maze spontaneous alternation test was used to assess the willingness of mice to explore new areas. The symmetrical Y-shaped Plexiglass maze had a white bottom and walls, and each mouse was allowed to move freely for 5 minutes. The observer recorded the time spent by the subject in each arm and manually recorded spontaneous alternations to ensure that the mouse explored a new arm each time it returned to the center of the maze. The percentage of spontaneous alternation was calculated as the number of consecutive entries into all three arms divided by the number of possible alternations. Novel object recognition test was conducted over three consecutive days. On the first day, mice explored the open field without any objects for 5 minutes. On the second day, two green-colored wooden cubes were introduced, and mice were allowed to explore for 5 minutes. On the third day, a novel object, an orange-colored wooden cylinder, was introduced along with a familiar object, a green-colored wooden cube. The mouse nose touching or pointing within 12 cm of the object was considered positive exploration, and the time spent exploring each object was manually recorded. The discrimination index was calculated using the ratio of time spent exploring the novel object to the time spent exploring both objects.

### Flow cytometry

Bone marrow cells were isolated from the tibia and femur by flushing a 25G syringe with 1X PBS containing 2% FBS. The red blood cells were removed using ACK lysis buffer before antibody incubation. For fetal liver cell analysis, timed pregnancies were performed, and embryos were harvested at different days post coitus into 1X PBS supplemented with 2% FBS. Fetal livers were isolated and dissociated using a pipette in fetal liver juice (FLJ), which consisted of 2% FBS, 2.5 mM EDTA, and 10 mM glucose in PBS. The cells were then counted and incubated with antibodies listed in the Supplemental Table 2. The cell populations were analyzed using BD Bioscience FACS Canto II, BD LSRII, and FACSymphony A5 flow cytometers.

### Single-cell RNA-sequencing and data analysis

The linage-marker-negative (Lin-) c-kit+ population containing hematopoietic stem and progenitor cells was sorted from dissociated mouse bone marrow cells by flow cytometry. Sorted single cell suspension, 10x barcoded gel beads, and oil (10xGenomics Chromium Next GEM Single Cell 3’ Kit v3.1 PN 1000268) were loaded into 10x Chromium™ Single Cell Chip to capture single cells into nanoliter-scale oil droplets by 10xChromium X Controller and generate Gel Bead-In-EMulsions (GEMs). cDNA from single cells were synthesized and barcoded by incubation of the GEMs in a thermocycler machine and then purified from GEMs by DynaBeads (10xGenomics PN 2000048). cDNA was pre-amplified by PCR to generate sufficient mass for sequencing library construction. The single cell cDNA libraries were constructed by following 10xGenomics 3’ NextGem 3.1 version kit instruction. The final constructed 3’-biased single cell libraries were sequenced by Illumina Novaseq6000 machine, targeting 50,000 read pairs/cell. Cellranger-7.1.0 (10X Genomics) pipeline was used to align the raw sequencing data to the mm10 mouse reference genome and count the expressed transcripts. Seurat (v4.3.0) was used to analyze and visualize the processed files. Detailed methods used for data analysis were described in Supplemental Methods.

## Statistics

Data are presented as ± SD. *P* values considered significant are presented below each figure. Statistical significance was determined by *t* test, one way ANOVA, non-parametric Kruskal-Wallis, and log-rank (Kaplan-Meier). Statistical analyses were performed using GraphPad Prism.

## Study approval

All the procedures of breeding and experiments using mice were approved and performed in accordance to the guidelines of the University of Alabama at Birmingham Institutional Animal Care and Use Committee.

## Data availability

Single-cell RNA-sequencing data will be deposited to publicly accessible databases upon acceptance of the manuscript for publication.

## Author contributions

LV, JHK, SSL, and EEA conceptualized the project. LV, BP, MM, HL, AE, ATD, JHK, KJ, JMM, RAB, AGS, RSW, BEH, SSL, and EEA designed the experiments, developed the methodology, and analyzed data. LV, BP, MM, HL, AE, ATD, JHK, KJ, JMM, LG, HGK, SNL, DS, and BEH performed the experiments and collected data. MM and EEA analyzed scRNA-Seq data, and JBF evaluated kidney H&E. LV, BP and EEA wrote the original draft of the manuscript with input from all authors. LV, BP, MM, AE, CAH, SD, LG, BEH, SSL, and EEA reviewed and edited the manuscript. EEA supervised the overall study.

## Acknowledgements

We thank Dr. Shanrun Liu and Vidya Sagar Hanumanthu at the University of Alabama at Birmingham (UAB) Flow Cytometry and Single Cell Core Facility for excellent technical support. We thank Dr. Bradley Yoder for evaluating the kidney histology slides, Dr. Maria Johnson for taking the bone µCT images, and Sharon for acquiring mouse brain MRI. We appreciate the support from the University of Alabama Health Services Foundation and UAB CF Research and Translation Core Center (P30DK072482) for the µCT instrument. This work was supported by the National Institutes of Health (NIH) grants (R01CA190688, R01CA236911, and R01HL168659 to E.E.A.; R01HL158800 and R01HL158875 to S.S.L.; K01OD026527 to B.E.H.), the institutional support from UAB School of Medicine, UAB Department of Pathology, and UAB O’Neal Comprehensive Cancer Center (to E.E.A.), and UAB Radiology Imaging Development Voucher program supported by the National Center for Advancing Translational Research/CCTS of the NIH under award number UL1TR001417.

## References

1. Tambuyzer E, Vandendriessche B, Austin CP, Brooks PJ, Larsson K, Miller Needleman KI, et al. Therapies for rare diseases: therapeutic modalities, progress and challenges ahead. Nat Rev Drug Discov. 2020;19(2):93–111.

2. Haendel M, Vasilevsky N, Unni D, Bologa C, Harris N, Rehm H, et al. How many rare diseases are there? Nat Rev Drug Discov. 2020;19(2):77–8.

3. Lee CE, Singleton KS, Wallin M, and Faundez V. Rare Genetic Diseases: Nature’s Experiments on Human Development. iScience. 2020;23(5):101123.

4. Farnaes L, Hildreth A, Sweeney NM, Clark MM, Chowdhury S, Nahas S, et al. Rapid whole-genome sequencing decreases infant morbidity and cost of hospitalization. NPJ Genom Med. 2018;3:10.

5. Hmeljak J, and Justice MJ. From gene to treatment: supporting rare disease translational research through model systems. Dis Model Mech. 2019;12(2).

6. Kim JH, Shinde DN, Reijnders MRF, Hauser NS, Belmonte RL, Wilson GR, et al. De Novo Mutations in SON Disrupt RNA Splicing of Genes Essential for Brain Development and Metabolism, Causing an Intellectual-Disability Syndrome. Am J Hum Genet. 2016;99(3):711–9.

7. Tokita MJ, Braxton AA, Shao Y, Lewis AM, Vincent M, Kury S, et al. De Novo Truncating Variants in SON Cause Intellectual Disability, Congenital Malformations, and Failure to Thrive. Am J Hum Genet. 2016;99(3):720–7.

8. Zhu X, Petrovski S, Xie P, Ruzzo EK, Lu YF, McSweeney KM, et al. Whole-exome sequencing in undiagnosed genetic diseases: interpreting 119 trios. Genet Med. 2015;17(10):774–81.

9. Takenouchi T, Miura K, Uehara T, Mizuno S, and Kosaki K. Establishing SON in 21q22.11 as a cause a new syndromic form of intellectual disability: Possible contribution to Braddock-Carey syndrome phenotype. Am J Med Genet A. 2016;170(10):2587–90.

10. Kim JH, Park EY, Chitayat D, Stachura DL, Schaper J, Lindstrom K, et al. SON haploinsufficiency causes impaired pre-mRNA splicing of CAKUT genes and heterogeneous renal phenotypes. Kidney Int. 2019;95(6):1494–504.

11. Dingemans AJM, Truijen KMG, Kim JH, Alacam Z, Faivre L, Collins KM, et al. Establishing the phenotypic spectrum of ZTTK syndrome by analysis of 52 individuals with variants in SON. Eur J Hum Genet. 2022;30(3):271–81.

12. Yang Y, Xu L, Yu Z, Huang H, and Yang L. Clinical and genetic analysis of ZTTK syndrome caused by SON heterozygous mutation c.394C>T. Mol Genet Genomic Med. 2019;7(11):e953.

13. Tan Y, Duan L, Yang K, Liu Q, Wang J, Dong Z, et al. A novel frameshift variant in SON causes Zhu-Tokita-Takenouchi-Kim Syndrome. J Clin Lab Anal. 2020;34(8):e23326.

14. Quintana Castanedo L, Sanchez Orta A, Maseda Pedrero R, Santos Simarro F, Palomares Bralo M, Feito Rodriguez M, et al. Skin and nails abnormalities in a patient with ZTTK syndrome and a de novo mutation in SON. Pediatr Dermatol. 2020;37(3):517–9.

15. Slezak R, Smigiel R, Rydzanicz M, Pollak A, Kosinska J, Stawinski P, et al. Phenotypic expansion in Zhu-Tokita-Takenouchi-Kim syndrome caused by de novo variants in the SON gene. Mol Genet Genomic Med. 2020;8(10):e1432.

16. Langford J, Vukadin L, Carey JC, Botto LD, Velinder M, Mao R, et al. SON-Related Zhu-Tokita-Takenouchi-Kim Syndrome With Recurrent Hemiplegic Migraine: Putative Role of PRRT2. Neurol Genet. 2023;9(3):e200062.

17. Ahn EY, DeKelver RC, Lo MC, Nguyen TA, Matsuura S, Boyapati A, et al. SON controls cell-cycle progression by coordinated regulation of RNA splicing. Mol Cell. 2011;42(2):185–98.

18. Sharma A, Takata H, Shibahara K, Bubulya A, and Bubulya PA. Son is essential for nuclear speckle organization and cell cycle progression. Mol Biol Cell. 2010;21(4):650–63.

19. Sharma A, Markey M, Torres-Munoz K, Varia S, Kadakia M, Bubulya A, et al. Son maintains accurate splicing for a subset of human pre-mRNAs. J Cell Sci. 2011;124(Pt 24):4286–98.

20. Hickey CJ, Kim JH, and Ahn EY. New discoveries of old SON: a link between RNA splicing and cancer. J Cell Biochem. 2014;115(2):224–31.

21. Lu X, Goke J, Sachs F, Jacques PE, Liang H, Feng B, et al. SON connects the splicing-regulatory network with pluripotency in human embryonic stem cells. Nat Cell Biol. 2013;15(10):1141–52.

22. Kim JH, Baddoo MC, Park EY, Stone JK, Park H, Butler TW, et al. SON and Its Alternatively Spliced Isoforms Control MLL Complex-Mediated H3K4me3 and Transcription of Leukemia-Associated Genes. Mol Cell. 2016;61(6):859–73.

23. Ilik IA, Malszycki M, Lubke AK, Schade C, Meierhofer D, and Aktas T. SON and SRRM2 are essential for nuclear speckle formation. Elife. 2020;9.

24. Alexander KA, Cote A, Nguyen SC, Zhang L, Gholamalamdari O, Agudelo-Garcia P, et al. p53 mediates target gene association with nuclear speckles for amplified RNA expression. Mol Cell. 2021;81(8):1666–81 e6.

25. Sun CT, Lo WY, Wang IH, Lo YH, Shiou SR, Lai CK, et al. Transcription repression of human hepatitis B virus genes by negative regulatory element-binding protein/SON. J Biol Chem. 2001;276(26):24059–67.

26. Ahn EY, Yan M, Malakhova OA, Lo MC, Boyapati A, Ommen HB, et al. Disruption of the NHR4 domain structure in AML1-ETO abrogates SON binding and promotes leukemogenesis. Proc Natl Acad Sci U S A. 2008;105(44):17103–8.

27. Candland DK, and Nagy ZM. The open field: some comparative data. Ann N Y Acad Sci. 1969;159(3):831–51.

28. Kraeuter AK, Guest PC, and Sarnyai Z. The Open Field Test for Measuring Locomotor Activity and Anxiety-Like Behavior. Methods Mol Biol. 2019;1916:99–103.

29. Simon P, Dupuis R, and Costentin J. Thigmotaxis as an index of anxiety in mice. Influence of dopaminergic transmissions. Behav Brain Res. 1994;61(1):59–64.

30. Kraeuter AK, Guest PC, and Sarnyai Z. The Y-Maze for Assessment of Spatial Working and Reference Memory in Mice. Methods Mol Biol. 2019;1916:105–11.

31. Lueptow LM. Novel Object Recognition Test for the Investigation of Learning and Memory in Mice. J Vis Exp. 2017(126).

32. Pietras EM, Reynaud D, Kang YA, Carlin D, Calero-Nieto FJ, Leavitt AD, et al. Functionally Distinct Subsets of Lineage-Biased Multipotent Progenitors Control Blood Production in Normal and Regenerative Conditions. Cell Stem Cell. 2015;17(1):35–46.

33. Iwasaki H, and Akashi K. Myeloid lineage commitment from the hematopoietic stem cell. Immunity. 2007;26(6):726–40.

34. Pronk CJ, Rossi DJ, Mansson R, Attema JL, Norddahl GL, Chan CK, et al. Elucidation of the phenotypic, functional, and molecular topography of a myeloerythroid progenitor cell hierarchy. Cell Stem Cell. 2007;1(4):428–42.

35. Hardy RR, and Hayakawa K. B cell development pathways. Annu Rev Immunol. 2001;19:595–621.

36. McKearn JP, Baum C, and Davie JM. Cell surface antigens expressed by subsets of pre-B cells and B cells. J Immunol. 1984;132(1):332–9.

37. Loder F, Mutschler B, Ray RJ, Paige CJ, Sideras P, Torres R, et al. B cell development in the spleen takes place in discrete steps and is determined by the quality of B cell receptor-derived signals. J Exp Med. 1999;190(1):75–89.

38. Purohit SJ, Stephan RP, Kim HG, Herrin BR, Gartland L, and Klug CA. Determination of lymphoid cell fate is dependent on the expression status of the IL-7 receptor. EMBO J. 2003;22(20):5511–21.

39. Mandel EM, and Grosschedl R. Transcription control of early B cell differentiation. Curr Opin Immunol. 2010;22(2):161–7.

40. Lukin K, Fields S, Hartley J, and Hagman J. Early B cell factor: Regulator of B lineage specification and commitment. Semin Immunol. 2008;20(4):221–7.

41. Inlay MA, Bhattacharya D, Sahoo D, Serwold T, Seita J, Karsunky H, et al. Ly6d marks the earliest stage of B-cell specification and identifies the branchpoint between B-cell and T-cell development. Genes Dev. 2009;23(20):2376–81.

42. Girschick HJ, Grammer AC, Nanki T, Mayo M, and Lipsky PE. RAG1 and RAG2 expression by B cell subsets from human tonsil and peripheral blood. J Immunol. 2001;166(1):377–86.

43. Sanchez M, Misulovin Z, Burkhardt AL, Mahajan S, Costa T, Franke R, et al. Signal transduction by immunoglobulin is mediated through Ig alpha and Ig beta. J Exp Med. 1993;178(3):1049–55.

44. Saleque S, Cameron S, and Orkin SH. The zinc-finger proto-oncogene Gfi-1b is essential for development of the erythroid and megakaryocytic lineages. Genes Dev. 2002;16(3):301–6.

45. Cantor AB, and Orkin SH. Transcriptional regulation of erythropoiesis: an affair involving multiple partners. Oncogene. 2002;21(21):3368–76.

46. Gerdes J, Lemke H, Baisch H, Wacker HH, Schwab U, and Stein H. Cell cycle analysis of a cell proliferation-associated human nuclear antigen defined by the monoclonal antibody Ki-67. J Immunol. 1984;133(4):1710–5.

47. Lu YC, Sanada C, Xavier-Ferrucio J, Wang L, Zhang PX, Grimes HL, et al. The Molecular Signature of Megakaryocyte-Erythroid Progenitors Reveals a Role for the Cell Cycle in Fate Specification. Cell Rep. 2018;25(8):2083–93 e4.

48. Ronzoni L, Bonara P, Rusconi D, Frugoni C, Libani I, and Cappellini MD. Erythroid differentiation and maturation from peripheral CD34+ cells in liquid culture: cellular and molecular characterization. Blood Cells Mol Dis. 2008;40(2):148–55.

49. Ohtsuka M, Inoko H, Kulski JK, and Yoshimura S. Major histocompatibility complex (Mhc) class Ib gene duplications, organization and expression patterns in mouse strain C57BL/6. BMC Genomics. 2008;9:178.

50. da Silva IL, Montero-Montero L, Ferreira E, and Quintanilla M. New Insights Into the Role of Qa-2 and HLA-G Non-classical MHC-I Complexes in Malignancy. Front Immunol. 2018;9:2894.

51. Dion W, Ballance H, Lee J, Pan Y, Irfan S, Edwards C, et al. Four-dimensional nuclear speckle phase separation dynamics regulate proteostasis. Sci Adv. 2022;8(1):eabl4150.

52. Rossi DJ, Bryder D, Zahn JM, Ahlenius H, Sonu R, Wagers AJ, et al. Cell intrinsic alterations underlie hematopoietic stem cell aging. Proc Natl Acad Sci U S A. 2005;102(26):9194–9.

53. Beerman I, Bhattacharya D, Zandi S, Sigvardsson M, Weissman IL, Bryder D, et al. Functionally distinct hematopoietic stem cells modulate hematopoietic lineage potential during aging by a mechanism of clonal expansion. Proc Natl Acad Sci U S A. 2010;107(12):5465–70.

54. Dykstra B, Olthof S, Schreuder J, Ritsema M, and de Haan G. Clonal analysis reveals multiple functional defects of aged murine hematopoietic stem cells. J Exp Med. 2011;208(13):2691–703.

55. Elias HK, Bryder D, and Park CY. Molecular mechanisms underlying lineage bias in aging hematopoiesis. Semin Hematol. 2017;54(1):4–11.

56. Henry CJ, Marusyk A, and DeGregori J. Aging-associated changes in hematopoiesis and leukemogenesis: what’s the connection? Aging (Albany NY). 2011;3(6):643–56.

57. Konturek-Ciesla A, Dhapola P, Zhang Q, Sawen P, Wan H, Karlsson G, et al. Temporal multimodal single-cell profiling of native hematopoiesis illuminates altered differentiation trajectories with age. Cell Rep. 2023;42(4):112304.

