## Supplemental for "A mouse model of ZTTK syndrome reveals indispensable SON functions in organ development and hematopoiesis"

### Supplemental Figures

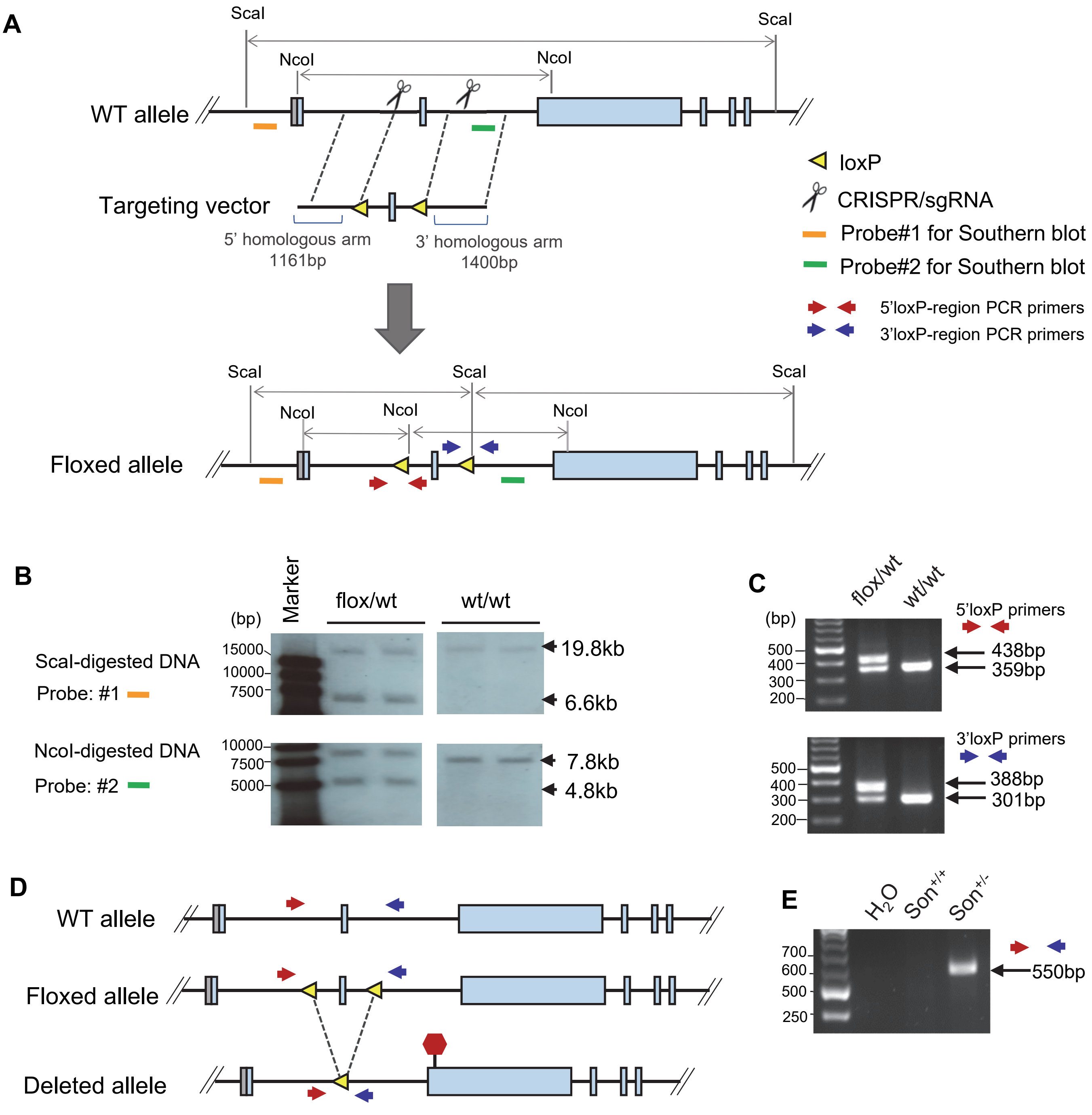


**Supplemental Figure 1. Generation of *Son*-floxed mice and *Son*$^{+/-}$ mice.** (**A**) Strategy for homologous recombination at the mouse *Son* gene locus and the locations of the probes and primers used for Southern blot and PCR shown in panels B, C and E. This design is based on ENSMUST00000114037.9 (mouse Son transcript-202 in Ensembl; NM_178880.4). (**B**) Southern blot analysis of ScaI-digested and NcoI-digested genomic DNA from tail snips of F1 mice that are heterozygous for the floxed allele, along with wild type control. (**C**) PCR genotyping to detect the floxed allele. (**D**) Schematics comparing the WT, floxed, and deleted allele and the location of a premature termination codon (red hexagon) formed upon deletion of the floxed region. (**E**) PCR genotyping to detect the deleted allele.

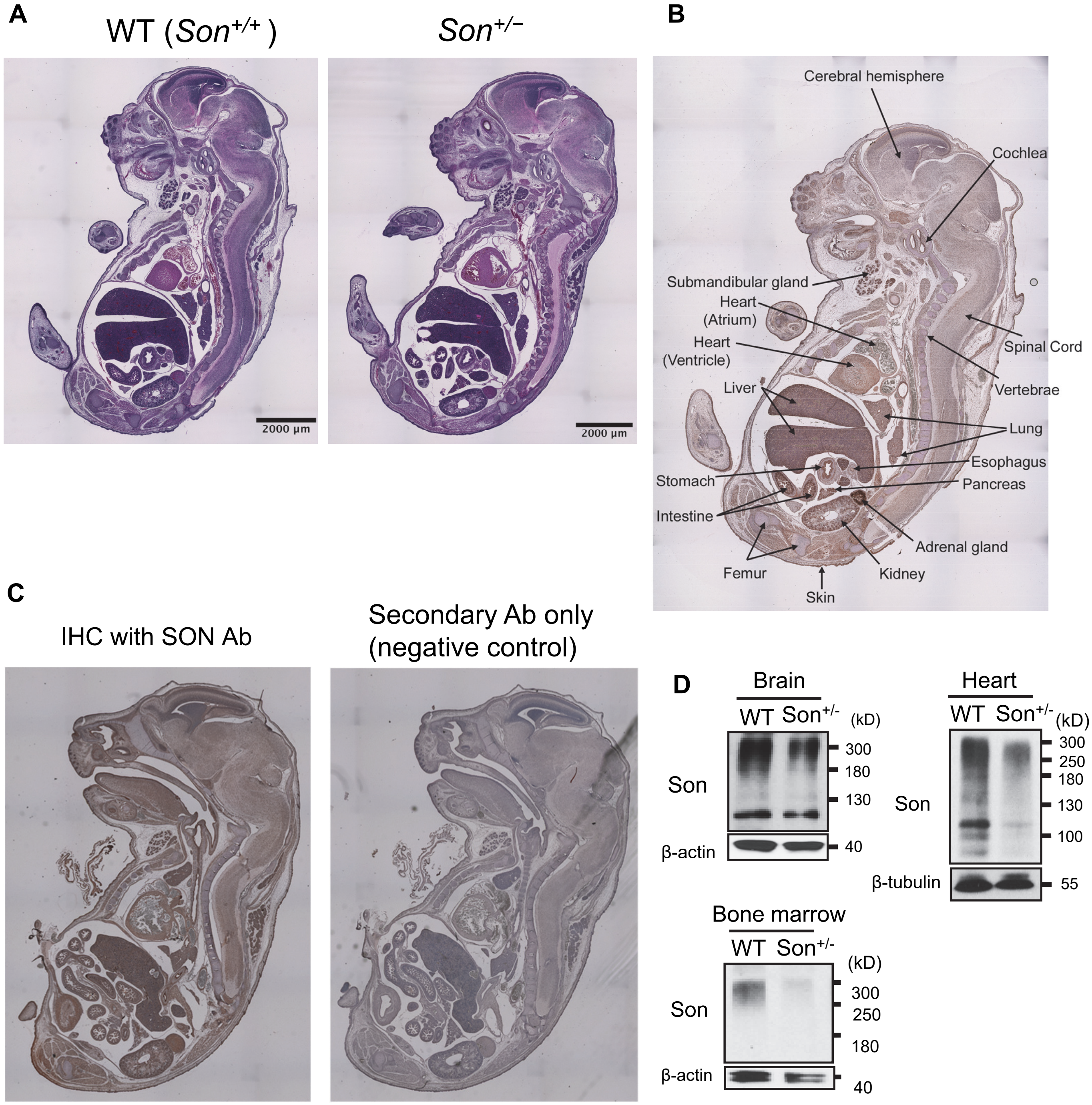


**Supplemental Figure 2. Analysis of Son expression in embryos and adult tissues of *Son*$^{+/-}$ mice.** (**A**) H&E staining of the sagittal section of WT and *Son*$^{+/-}$ embryos (E16), showing no significant defects in forming fetal organs in the *Son*$^{+/-}$ embryos. (**B**) An image of an E16 WT embryo and annotations of tissue/organs.(**C**) IHC staining without primary antibody (secondary antibody only) is completely devoid of immunopositivity, indicating specificity of brown staining detected by Son antibody (Ab). (**D**) The expression of Son in the indicated organs of WT and *Son*$^{+/-}$ adult mice was analyzed by Western blot.

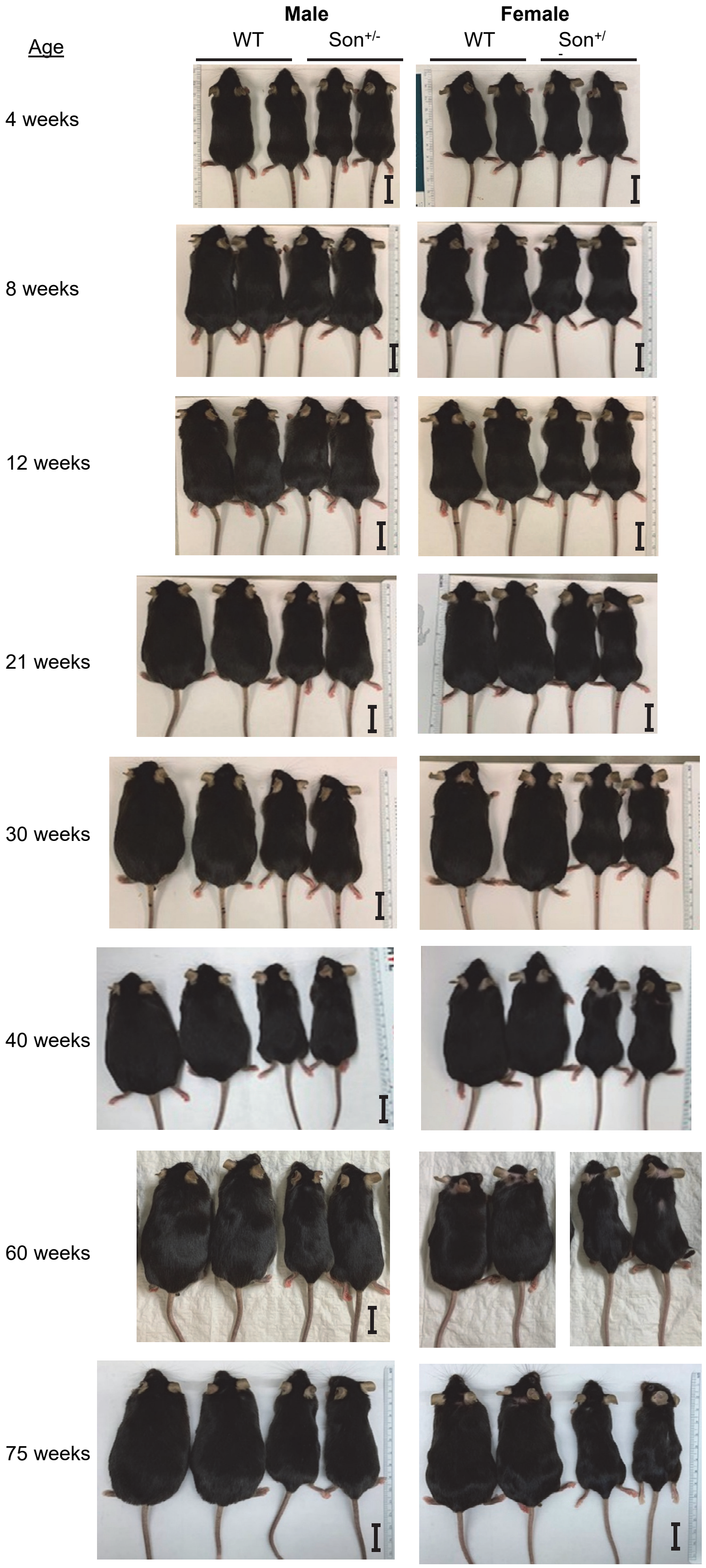


2 cm

| Age | | Male WT | Male Son+/- | Male p | Female WT | Female Son+/- | Female p |
|---|---|---|---|---|---|---|---|
| 4 weeks | Weight (g) | 18.8 ± 0.8 | 16.3 ± 2.1 | $p < 0.01$ | 16.7 ± 0.6 | 13.0 ± 1.0 | $p < 0.01$ |
| | Length (cm) | 8.5 ± 0.1 | 8.0 ± 0.0 | $p < 0.001$ | 8.3 ± 0.3 | 8.0 ± 0.0 | $p = 0.1161$ |
| 8 weeks | Weight (g) | 25.0 ± 0.7 | 21.5 ± 0.8 | $p < 0.001$ | 20.7 ± 1.2 | 16.3 ± 0.6 | $p < 0.01$ |
| | Length (cm) | 9.7 ± 0.2 | 9.0 ± 0.1 | $p < 0.01$ | 9.1 ± 0.2 | 8.7 ± 0.1 | $p < 0.05$ |
| 12 weeks | Weight (g) | 29.6 ± 0.5 | 23.0 ± 1.7 | $p < 0.001$ | 24.0 ± 1.7 | 18.3 ± 0.6 | $p < 0.01$ |
| | Length (cm) | 10.0 ± 0.0 | 9.4 ± 0.3 | $p < 0.05$ | 9.5 ± 0.5 | 9.0 ± 0.0 | $p = 0.1583$ |
| 21 weeks | Weight (g) | 39.8 ± 1.3 | 27.5 ± 2.9 | $p < 0.001$ | 32.0 ± 3.6 | 20.3 ± 1.5 | $p < 0.01$ |
| | Length (cm) | 10.7 ± 0.1 | 9.7 ± 0.1 | $p < 0.001$ | 10.2 ± 0.1 | 18.9 ± 0.1 | $p < 0.001$ |
| 30 weeks | Weight (g) | 44.3 ± 2.3 | 27.5 ± 2.3 | $p < 0.001$ | 35.2 ± 4.4 | 20.9 ± 1.7 | $p < 0.01$ |
| | Length (cm) | 10.8 ± 0.2 | 9.8 ± 0.2 | $p < 0.001$ | 10.2 ± 0.1 | 9.2 ± 0.3 | $p < 0.01$ |
| 40 weeks | Weight (g) | 47.7 ± 3.3 | 30.0 ± 1.7 | $p < 0.001$ | 37.8 ± 5.5 | 21.6 ± 0.9 | $p < 0.01$ |
| | Length (cm) | 11.0 ± 0.1 | 9.7 ± 0.1 | $p < 0.001$ | 10.4 ± 0.1 | 9.1 ± 0.1 | $p < 0.001$ |
| 60 weeks | Weight (g) | 53.0 ± 2.5 | 30.8 ± 1.1 | $p < 0.001$ | 39.3 ± 4.3 | 22.8 ± 1.3 | $p < 0.05$ |
| | Length (cm) | 10.6 ± 0.1 | 9.5 ± 0.2 | $p < 0.001$ | 10.6 ± 0.1 | 9.3 ± 0.3 | $p < 0.01$ |

**Supplemental Figure 3. Photos of male and female *Son*$^{+/-}$ mice at different ages with gender-matched WT littermates at different ages.** Next to the photos, the body weight and body length were indicated as average ± SD for each group. n=3-5 per group, p values from two-tailed, unpaired t-test were indicated for each group. Scale bar, 2 cm.

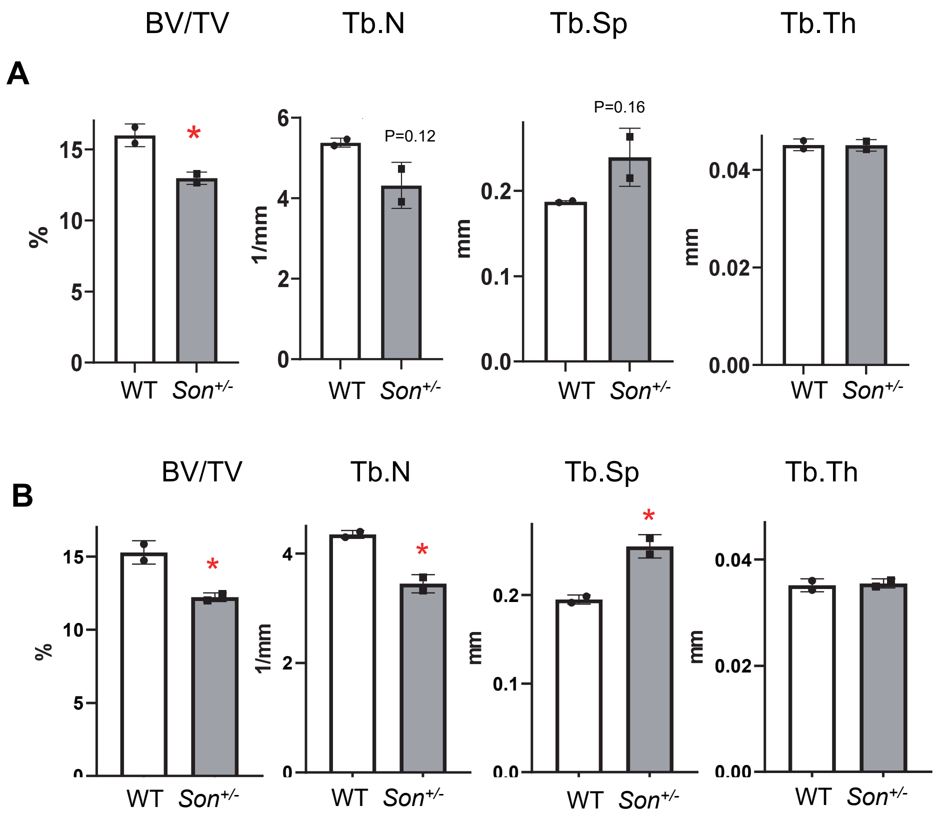


**Supplemental Figure 4. Morphometric analyses of microCT data of femurs from 6-week-old female *Son*$^{+/-}$ and control mice using the TRI plate model of analysis.** microCT Scans were reconstructed into 2-D slices and all slices analyzed using the µCT Evaluation Program (v6.5-2, Scanco Medical) and 3-D reconstruction performed using µCT Ray v4.2. Trabecular bone was evaluated in the distal femoral metaphysis (density threshold of 501 mg HA/cm3). For trabecular bone, the bone volume fraction (BV/TV), trabecular number (Tb.N), trabecular spacing (Tp.Sp), and trabecular thickness (Tb.Th) were determined using the DT analysis method (**A**) and TRI model analysis (**B**), according to the protocol https://www.scanco.ch/faq.html. n=2 per genotype, *$p < 0.05$.

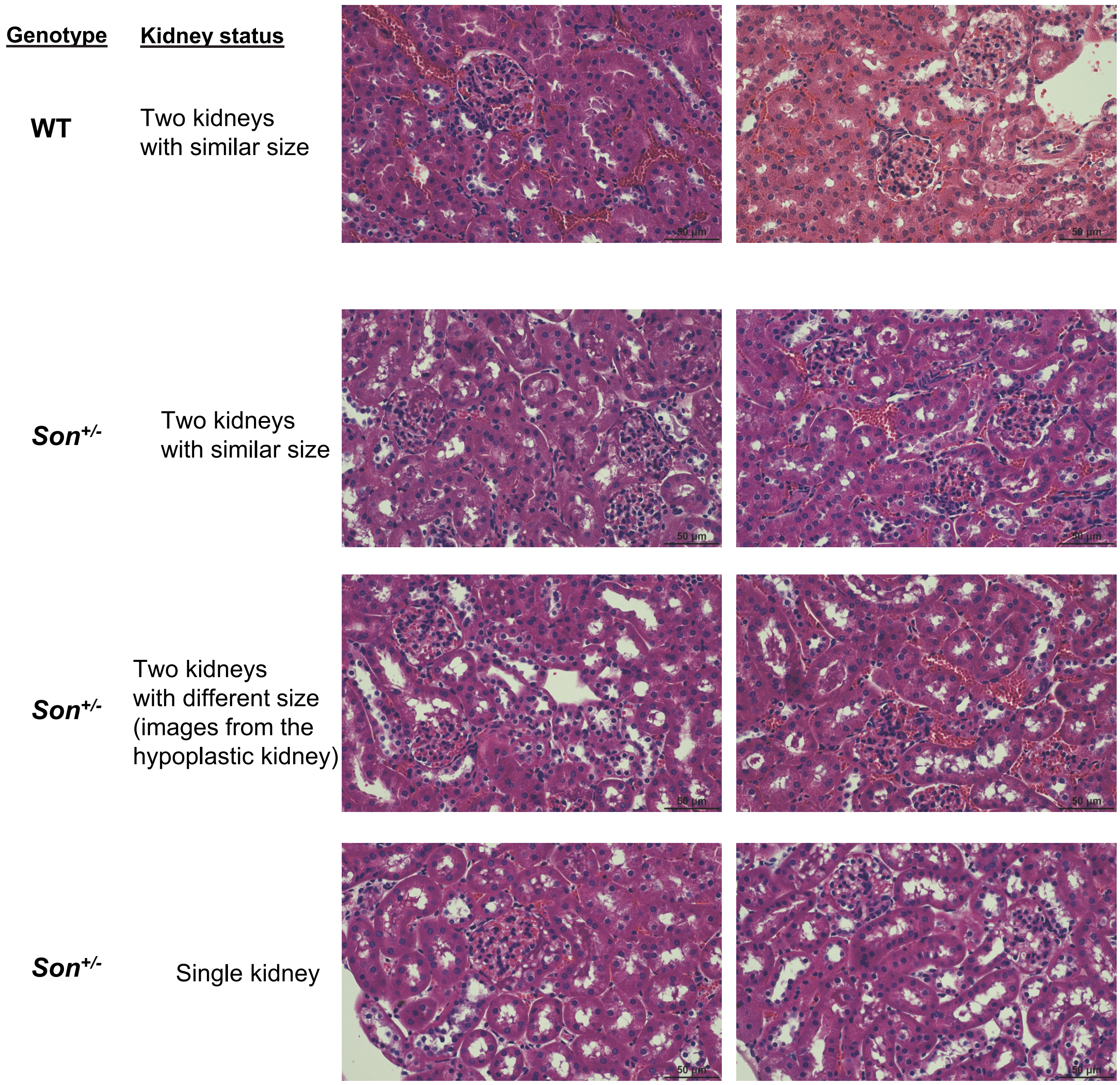


**Supplemental Figure 5. H&E staining of the kidney sections from WT and *Son*$^{+/-}$ mice with indicated kidney status.** Mild tubular regeneration and interstitial edema were observed in Son+/- kidneys, especially in the sample from Son+/- mice with a single kidney.

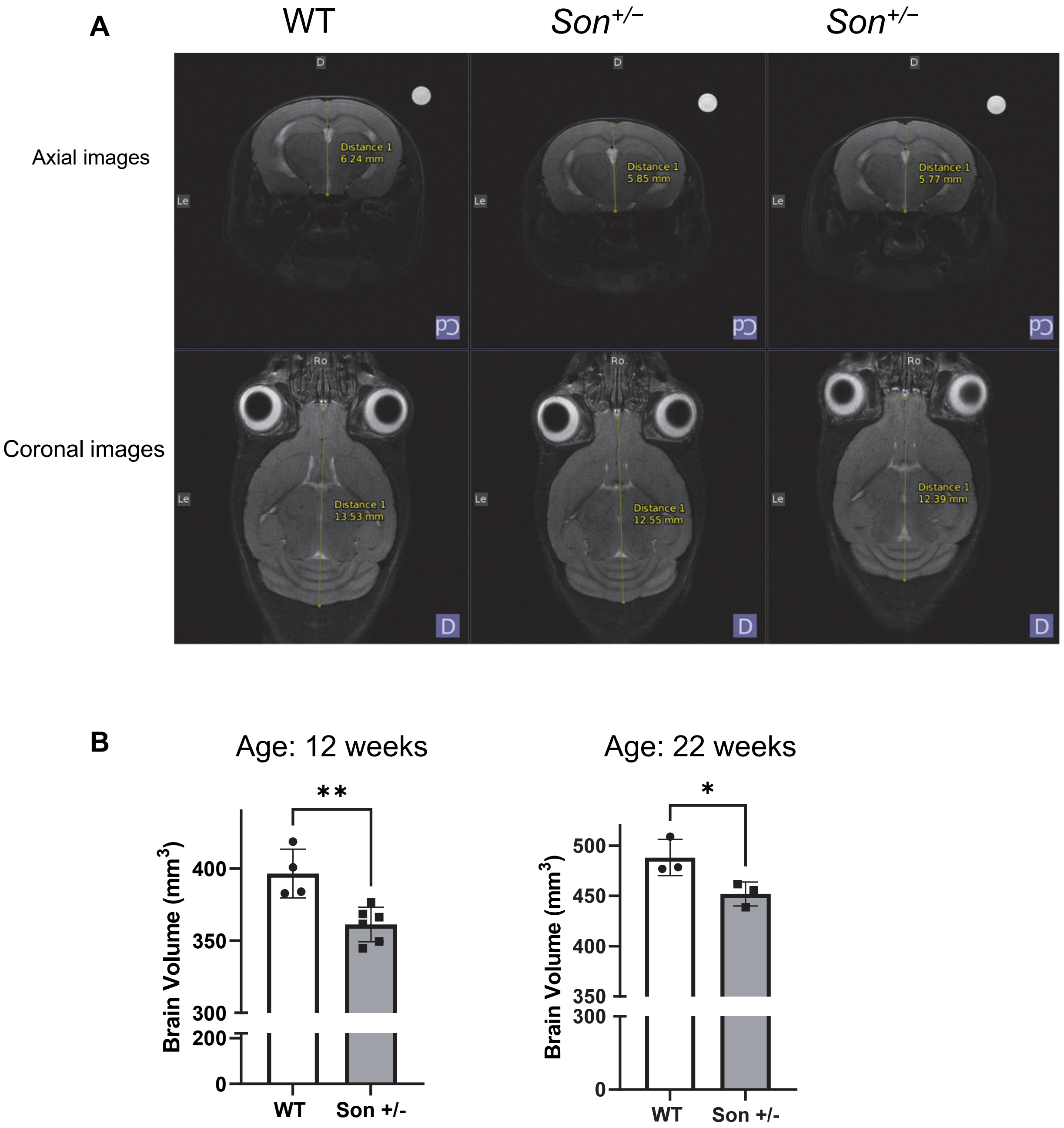


**Supplemental Figure 6. Magnetic resonance imaging (MRI) of the brain indicates that *Son*$^{+/-}$ mice have smaller brain volumes compared to WT mice of matched age**. (**A**) Representative T2-weighted brain MRI images of 10-week-old WT and *Son*$^{+/-}$ mice. MRI was performed using a preclinical 9.4T MRI scanner (Bruker BioSpin MRI GmbH, Ettlingen, Germany) with an 86 mm inner-diameter quadrature volume coil for excitation and a 2x2 brain surface coil array for signal reception. (**B**) Graphs indicating the brain volumes calculated from the MRI data. High-resolution T2-weighted images were acquired using a multi-slice rapid acquisition with relaxation enhancement (RARE) spin-echo pulse sequence with the following parameters: TR/TE = 2000 ms / 40 ms; NEX = 8; RARE factor = 8; bandwidth = 66.7 kHz; FOV = (20 x 20) mm; matrix size = minimum of (256 x 256); slice thickness = 0.6-0.7 mm. Regions of interest (ROIs) were drawn around the entirety of the brain in each image slice using ImageJ Fiji. The area of each ROI was calculated using ImageJ Fiji and multiplied by the image slice thickness to yield an estimated volume measurement for the entire brain. Mean ± SD, * $p < 0.05$, ** $p < 0.01$

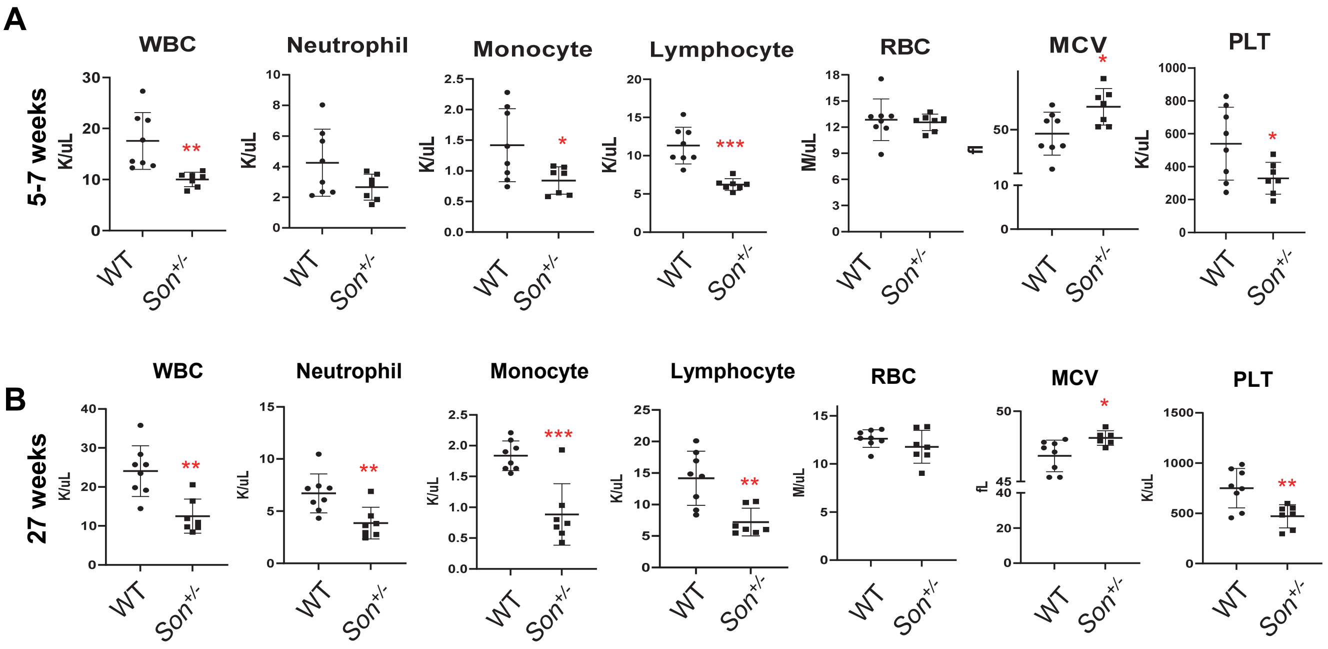


**Supplemental Figure 7. Complete blood counts (CBC) from whole blood of the *Son*$^{+/-}$ mice showed the abnormalities which are similar to those observed in human ZTTK syndrome patients.** CBC were performed for *Son*$^{+/-}$ mice and WT littermates at age 5-7 weeks (**A**) and 27 weeks. (**B**) Persistent abnormalities found in *Son*$^{+/-}$ mice include decrease of overall white blood cells (WBCs), neutrophils, monocytes, lymphocytes, and platelets. Red blood cell (RBC) counts are in normal range, but MCV is significantly increased in *Son*$^{+/-}$ mice, indicating impaired terminal differentiation of RBCs. Data are presented as mean ± SD, n=7-8.

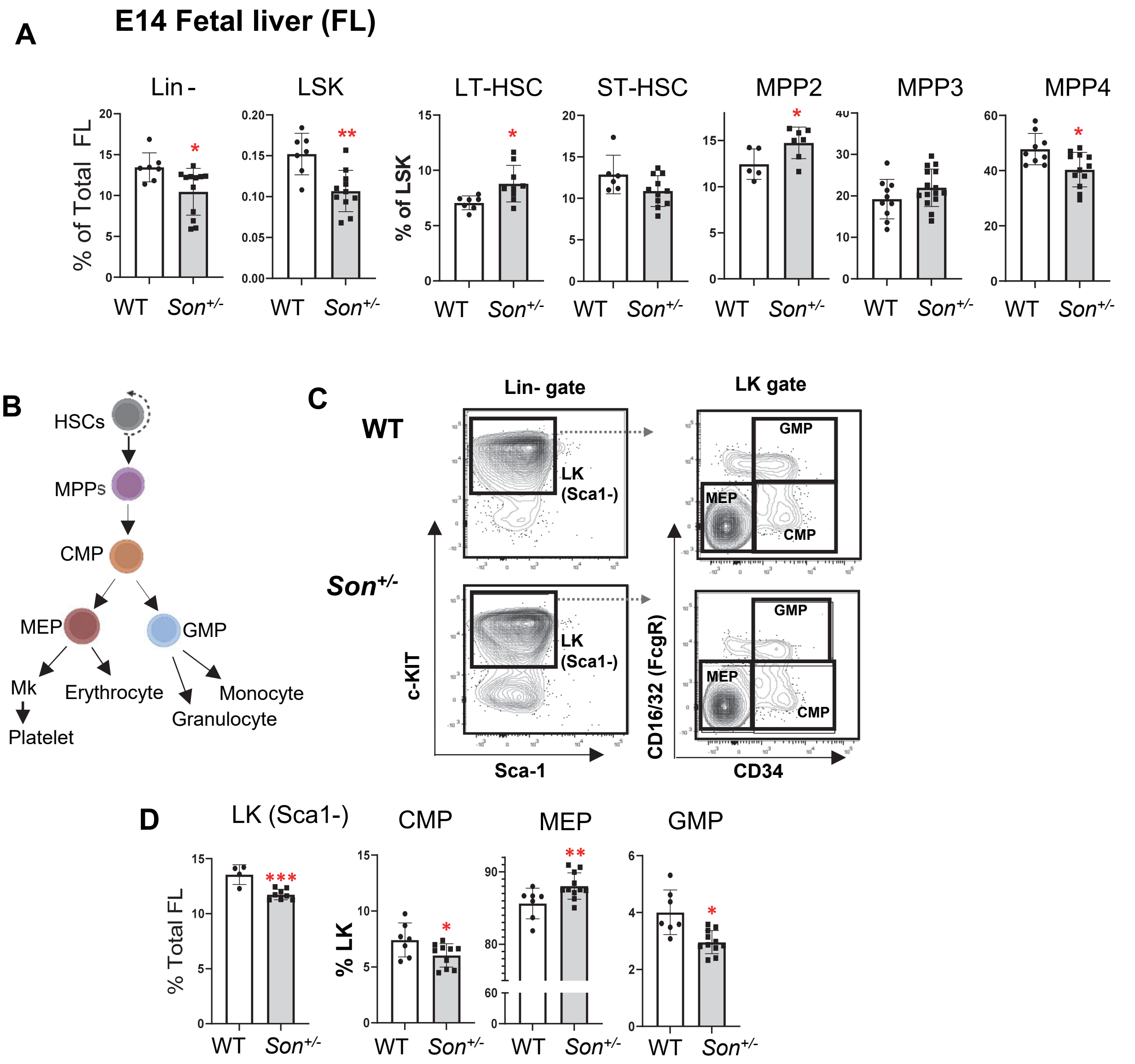


**Supplemental Figure 8. Analyses of fetal liver hematopoiesis identified altered HSPC subpopulations and an imbalance of myeloid progenitor lineage fate in *Son*$^{+/-}$ embryo.** (**A**) Frequency of the indicated populations within total fetal liver cells from E14 embryos (for Lin- and LSK) or within the LSK population (LT-HSC, ST_HSC, MPP2, MPP3 and MPP4). Data are expressed as mean ± SD, n=8-11, *p < 0.05, **p < 0.01, ***p < 0.001. (**B**) A schematic depicting the classical model of myeloid progenitor differentiation. (**C**) Flow cytometry contour plots showing the gating scheme for LK(Sca1-), CMP, GMP and MEP. (**D**) Frequency of indicated populations within E14 fetal liver cells (for LK, Sca1-) or within the LK population (for CMP, MEP, and GMP). Data are expressed as mean ± SD, n=5-11, *p < 0.05, **p < 0.01, ***p < 0.001.

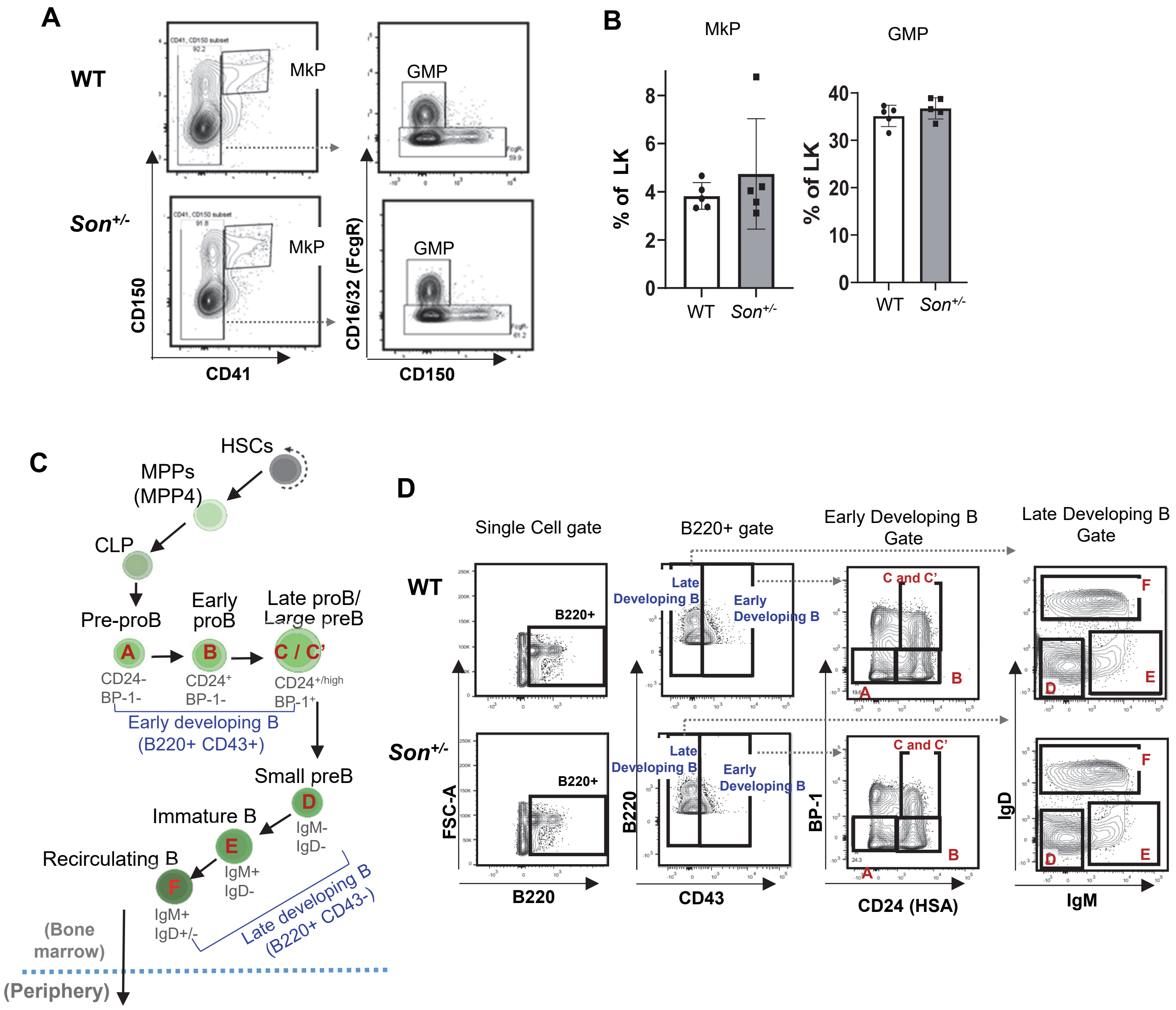


**Supplemental Figure 9. Analyses of myeloid progenitors and B cell progenitor in the bone marrow Schematics of bone marrow B cell analysis.** (**A**) Flow cytometry contour plots showing the gating scheme for MkP (megakaryocyte progenitors) and GMP (granulocyte/monocyte progenitors). (**B**) Frequency of MkP and GMP within bone marrow LK (Lin- cKit+) cells, expressed as mean ± SD, n=5. (**C**) A schematic depicting the stages of B cell development in the bone marrow. Hardy fractions A – F (in red font, based on Hardy et al. 1991) ) are shown with surface markers,. (**D**) Flow cytometry contour plots showing the gating scheme for Hardy fractions A – F, by determining the expression status of B220, CD43, CD24, BP-1, IgM, and IgD.

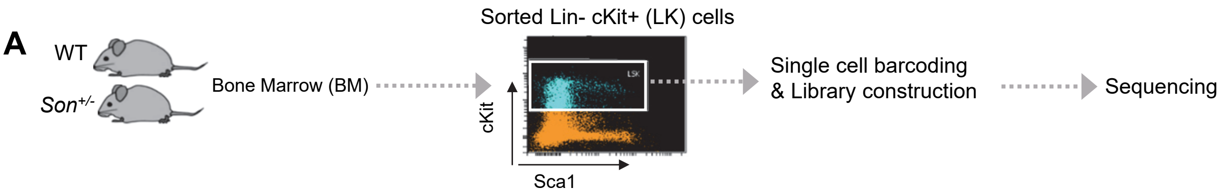


**B**

| Sample | Number of analyzed cells | Number of annotated cells |
|---|---|---|
| $Son^{+/-}$-1 | 11,887 | 11,764 |
| $Son^{+/-}$-2 | 19,634 | 19,468 |
| WT-1 | 12,264 | 12,173 |
| WT-2 | 12,275 | 12,182 |
| $Son^{+/-}$ Total | 31,521 | 31,232 |
| WT Total | 24,539 | 24,355 |

Filteration criteria:

- •nUMI >= 500
- •nGene >= 250
- •log10GenesPerUMI > 0.80
- •mitoRatio < 0.20

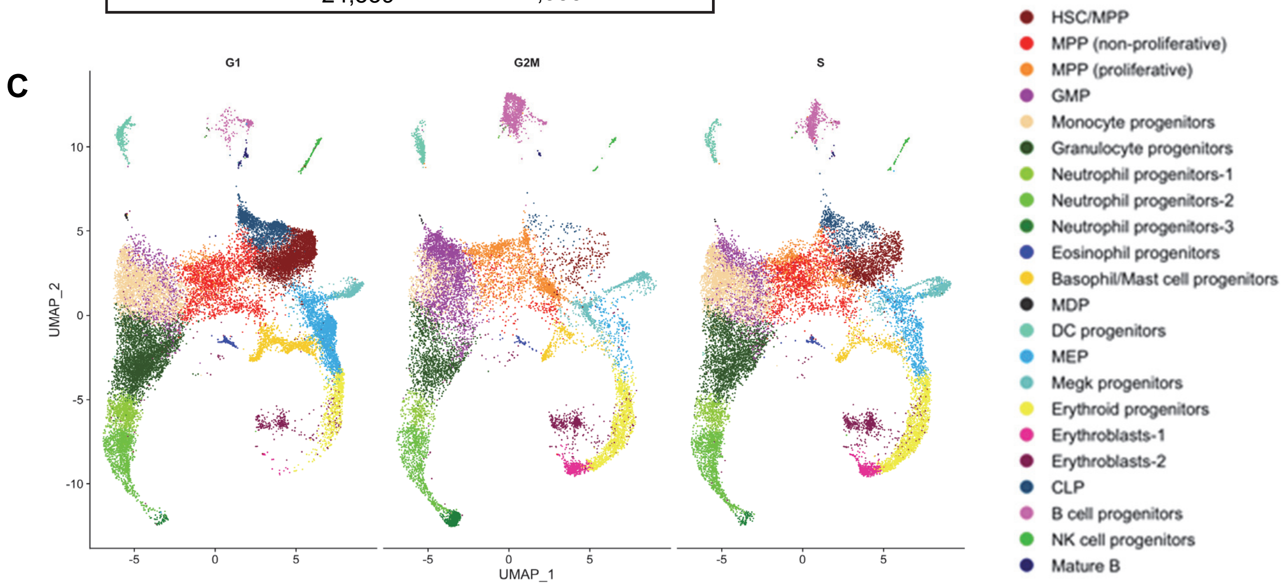


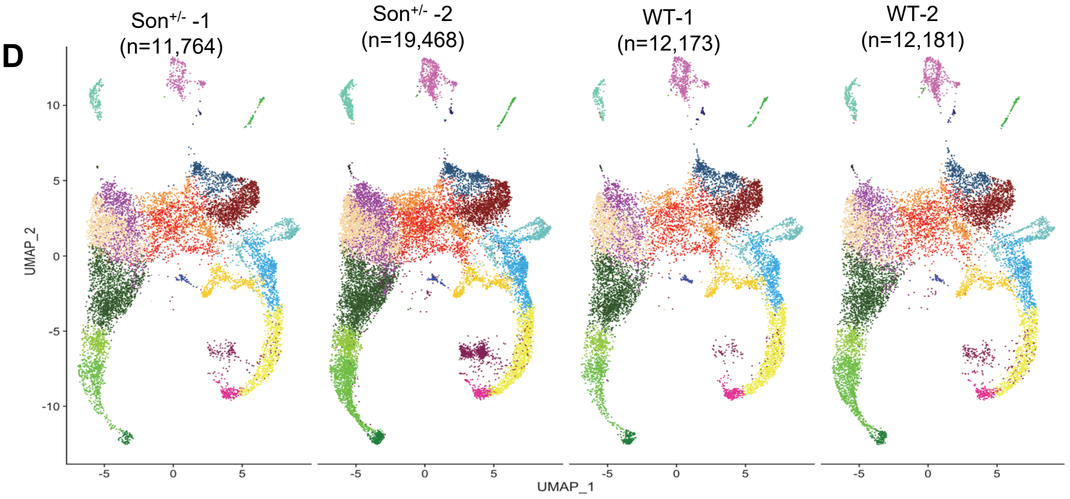


**Supplemental Figure 10. Quality control, filtering criteria and UMAP lots of single-cell RNA-sequencing (scRNAseq) data.** (**A**) Schematics of the scRNAseq procedure. (**B**) Numbers of cells analyzed and annotated after filtration using the criteria listed. (**C**) Cell cycle status (G1, G2/M, and S phases) of the 22 clusters (color-coded) identified by Seurat. (**D**) UMAP plots showing the 22 clusters identified in the bone marrow LK cells from WT and $Son^{+/-}$ mice. The numbers of annotated/plotted cells per each sample are indicated.

**Supplemental Table 1. Hematological abnormalities identified in human ZTTK syndrome patients*.**

| | | |
|---|---|---|
| **Myeloid lineage phenotype** | **Megakaryocyte / Erythroid lineage** | • Large RBC (High MCV)<br>• High and Low RBC<br>• Polycythemia<br>• Thalassemia (Low Hemoglobin)<br>• Low Platelets<br>• Deep Vein Thrombosis (DVT) (Blood clots, Stroke) |
| | **Granulocyte / Monocyte lineage** | • High and Low WBC<br>• High and Low Neutrophils |
| **Lymphoid lineage phenotype** | | • Low lymphocyte counts<br>• Immunoglobulin deficiency (IgG, IgA, and IgM)<br>• Poor response to vaccine<br>• Recurrent Infection (ex. respiratory infection, severe allergy, severe reactions to common illnesses, common variable immune deficiency (CVID))<br>• Intravenous Immunoglobulin (IVIG) monthly<br>• Autoimmune disorders (ex. Autoimmune encephalitis) |
| **Others** | | • Bone marrow failure<br>• High Vit. B12 in serum<br>• Low Vit. A and Vit. E |

*Voluntary reports from patients and families to the ZTTK SON-Shine Foundation, https://zttksonshinefoundation.org/

**Supplemental Table 2. Antibodies used for flow cytometry analysis and sorting.**

| Antibodies (anti-Mouse) | Source | Identifier |
|---|---|---|
| Lineage antibody-**FITC** (clones: CD3 (17A2), CD11b (M1/70), CD45/B220 (RA3-6B2), Ly-6G (Gr-1) (RB6-8C5) and TER-119 (TER-119) | Tonbo Bioscience | Fisher catalog # 50-105-5256 |
| CD117 (c-Kit)-**APC** (clone 2B8) | BD Biosciences | BD catalog # BD553356 |
| Sca-1-**PerCP-cy5.5** (clone D7) | Biolegend | Biolegend catalog # 108123 |
| CD135-**PE** (clone A2F10) | Biolegend | Biolegend catalog # 135305 |
| CD150-**PE-cy7** (clone mShad150) | Invitrogen | Fisher catalog #50-245-735 |
| CD48-**APC-Cy7** (clone HM48-1) | BD Biosciences | BD catalog # BD561242 |
| Live/Dead 405/545 stain (**BV510**) | Biotium | Fisher catalog # NC1907443 |
| Sca-1-**BV421** (clone E13-161.7) | Biolegend | Biolegend catalog #108123 |
| CD16/32 (FcRII)-**APC-cy7** (clone 93) | Biolegend | Biolegend catalog # 101327 |
| CD41-**BV711** (clone MWReg30) | BD Biosciences | BD catalog # BDB740712 |
| CD71-**PE** (clone C2) | BD Biosciences | BD catalog # BDB553267 |
| CD105 (Endoglin)-**BV605** (clone MJ7/18) | BD Biosciences | BD catalog # BDB740425 |
| Ter-119-**PE-cy5** (clone TER-119) | Invitrogen | Fisher catalog #15592181 |
| CD45R/B220-**APC-cy7** (clone RA3-6B2) | Biolegend | Fisher catalog # NC9009216 |
| CD3-**APC-cy7** (clone 17A2) | BD Biosciences | BD catalog #BDB560590 |
| CD45R/B220-**Pacific Blue** (clone RA3-6B2) | BD Biosciences | BD catalog #BDB558108 |
| CD11b-**Pacific Blue** (clone M1/70) | Southern Biotech | Fisher catalog # OB156126 |
| Gr-1-**Pacific Blue** (clone RB6-8C5) | Biolegend | Fisher catalog # 50-163-264 |
| BP-1/Ly-51-Biotin (clone 6C3) | Biolegend | Fisher catalog # 108303 |
| Streptavidin-**V500** | BD Biosciences | BD catalog # BDB561419 |
| CD43-**FITC** (clone S7) | BD Biosciences | BD catalog # BDB553270 |
| CD24-**PE** (clone M1/69) | BD Biosciences | BD catalog #BDB553262 |
| CD23-**PE-Cy7** (clone B3B4) | eBioscience | Fisher catalog # 50-154-79 |
| IgD-**PerCP-eFluor 710** (clone 11-26c (11-26)) | eBioscience | Fisher catalog # 50-112-8623 |
| IgM-**APC** (clone II/41) | eBioscience | Fisher catalog # 50-152-13 |
| CD19-**APC-Cy7** (clone 1D3) | BD Biosciences | BD catalog # BDB557655 |
| CD21-**FITC** (clone 7G6) | BD Pharminogen | Fisher catalog #561769 |
| CD93-**PE** (clone AA4.1) | Invitrogen | Fisher catalog #12-5892-81 |

### Supplemental Methods

#### Genotyping

Tail or ear snips were incubated in 113 µL 50 mM NaOH at 95°C for 30 minutes, followed by adding 32 µL of 1M Tris-buffer (pH 8.8) and centrifugation for 10 minutes. Supernatant was used for the following PCR reaction: 95°C for 5 minutes initiation, 95°C for 30 seconds, 58.5°C for 30 seconds, 72°C 1 kb/min for 35 cycles for elongation, and 72°C for 10 minutes for termination. Primer sequences to detect the presence of the LoxP sequence are the following: 3' LoxP site (Forward: AGAACATGGCCACCCATTTCTTCCA, Reverse: ATCCCTCTCTAGGAGTTGCTGGTGT) and 5' LoxP site (Forward: TGCCACTTGTTCTGTGTAAAATTCTTGA, Reverse: GCCACACTGTGCACTGATCACAAAC). To detect the deletion of the LoxP-floxed sequence, the forward primer for 5' LoxP and the reverse primer for 3' LoxP listed above were used.

#### Analysis of single-cell RNA-sequencing data

Cellranger-7.1.0 (10X Genomics) pipeline was used to align the raw sequencing data to the mm10 mouse reference genome and count the expressed transcripts. Seurat (v4.3.0) was used to analyze and visualize the processed files. For quality control and filtration, cells were filtered using four criteria; (a) the gene number per cell to be more than 250, (b) the UMI counts of more than 500 per cell, (c) log10 genes per UMI of more than 0.80, and (d) mitochondrial gene expression per cell less than 20% of total gene expression. Normalization and variance stabilization of filtered row counts was done using the SCtransform method to account for the variation due to sequencing depth.

To ensure that the cell types of wild-type align with the same cell types of knockout, samples were integrated using shared highly variable genes from each condition. The top 3000 highly variable features were selected using *SelectIntegrationFeatures* and *PrepSCTIntegration* functions. They have been used to perform canonical correlation analysis (CCA) to correct the batch effect and to find anchors between two datasets using the mutual nearest neighbors (MNNs) algorithm using *FindIntegrationAnchors*. The final integration step was done by the *IntegrateData* function.

For clustering analysis and annotation, a graph-based clustering method using a K-nearest neighbor algorithm (*FindNeighbors* function) was applied based on the top 40 PCA, then performing the clustering step using the *FindClusters* function to group cells together with a resolution of 0.6. Cell clusters were visualized using UMAP by first applying the *RunPCA* function, then *RunUMAP* with dims parameter 1:40. Clusters annotation was done manually based on the conserved marker genes identified using the *FindConservedMarkers* function. Then, these genes were used in the mouse hematopoietic marker gene sets database (CellKb Immune v2.3) to label the clusters.
